# DPP9-mediated inflammasome repression protects against checkpoint inhibitor lung toxicity

**DOI:** 10.64898/2026.06.30.735609

**Authors:** J. Richard Brewer, Ailin Han, Amin H. Nassar, Elias Bou Farhat, Holly N. Blackburn, Tianli Xiao, Haris Mirza, Walter K. Mowel, Esen Sefik, Saskia Hartner, Michael Chiorazzi, Takeshi Ito, Min-Hee Oh, Matthew Z. Madden, Athreya Rangavajhula, Elio Adib, Mustafa J. Saleh, Marc Machaalani, Mehrdad Rakaee, Masoud Tafavvoghi, Claire Quattropani, Nicole Gazetos, David Gerber, Farjana Fattah, Jeffrey A. SoRelle, Dominic Choo, Mitchell S. von Itzstein, Shannon Bevans-Fonti, Mohammad Ghanbar, Karthik Suresh, Thomas Mazumder, Chun J. Ye, Toni K. Choueiri, Alexander Gusev, Richard A. Flavell

**Author notes:** These authors contributed equally to this work (J. Richard Brewer, Ailin Han, Amin H. Nassar).

## Abstract

Over one million patients receive cancer immunotherapy annually, yet the mechanisms underlying life-threatening immune-mediated toxicities remain poorly understood. Checkpoint inhibitor pneumonitis (CIP) is the leading cause of immunotherapy-related mortality, with a case fatality rate approaching 10%, and no genetic risk factors have been described to date. We identified Dipeptidyl-peptidase 9 (DPP9) as the first genetic susceptibility gene for CIP in a clinico-genomics cohort of 4,397 patients treated with immune checkpoint inhibitors. Mechanistically, DPP9 suppresses CARD8 inflammasome activation and IL-18 secretion in human monocytes, a pathway which is engaged prior to CIP onset, with IL-18 selectively elevated in the plasma of patients who subsequently develop CIP. Myeloid-restricted ablation of *Dpp8* and *Dpp9* in mice recapitulated the pulmonary histopathological and immunological hallmarks of CIP, including granuloma formation, accumulation of IFNγ-producing T cells and monocyte-derived macrophages. Each of these phenotypes were driven by excessive IL-18 secretion. Together, these findings establish DPP9 as a genetic determinant of CIP and nominate IL-18 blockade as a mechanistically rational therapeutic strategy.

## Introduction

In recent years, immune checkpoint inhibitor (ICI) therapy has transformed cancer treatment, leading to dramatic improvements in survival in many cancer types^1,2^. However, this broad application of ICI therapy has been accompanied by a concomitant rise in therapy limiting immune-related adverse events across multiple organ systems. Checkpoint inhibitor pneumonitis (CIP) is characterized by widespread inflammation of the lungs and is among the most clinically significant toxicities^3^. Real-world data suggests a high prevalence of CIP, ranging from 10% to 20%^3–5^. Indeed, CIP is the leading cause of mortality related to ICI therapy^3,6,7^. Survivors of CIP have limited treatment options and often worse overall survival^4,8^. Chronic inflammation caused by CIP can result in pulmonary fibrosis, further reducing survival odds. Understanding the heritable risk and pathways that regulate both inflammatory and restorative responses in CIP is essential for risk-stratifying patients and developing targeted strategies to minimize tissue damage, promote tissue repair, and prevent fibrosis.

Emerging studies have begun to describe the cellular underpinnings of CIP in the clinic^9,10^. These investigations have uncovered significant perturbations in immune cell composition and intercellular dynamics in patients with CIP, including: (i) increased frequency of TNF-α^high^, IFN-γ^high^ CD8^+^ T cells, (ii) fewer suppressive regulatory T cells, (iii) an increased presence of central memory T cells, (iv) the emergence of pathogenic T-helper 17.1 cells characterized by the expression of TBX21, RORC, IFNγ, IL-17A, and CSF2, and (v) altered myeloid cell composition with fewer alveolar macrophages, and more monocyte-derived macrophages/dendritic cells (DCs) ^9–12^. Although our prior genome-wide association study (GWAS) has identified specific single nucleotide polymorphisms (SNPs) near IL-7 associated with a heightened risk of any grade immune-related adverse events, these have not been explicitly linked to discrete immune-related adverse events including CIP^13^. As such, genetic association studies to date in this domain have been constrained by limited cohort sizes, the conflation of disparate toxicities, and the challenges of accurately curating toxicity data from real-world settings.

To identify genes that influence CIP development, we considered risk factors for other inflammatory lung disorders. GWAS have demonstrated that susceptibility to SARS-CoV-2 infection and idiopathic pulmonary fibrosis (IPF) are heritable and associated with specific genetic variants^14–18^. GWAS have identified three intronic DPP9 SNPs among the most robustly replicated risk variants for both severe SARS-CoV-2 infection^14–16^ and IPF^17,18^. These associations between *DPP9* variants and severe COVID-19 have also been validated across diverse ancestral populations^16^. DPP9, and closely related DPP8, are peptidases that serve as critical negative regulators for the cytoplasmic pathogen sensors CARD8 and NLRP1^19–22^. Upon activation, CARD8 and NLRP1 initiate an oligomerization-based signaling cascade that promotes the recruitment of CASP1 into a supramolecular protein complex known as the “inflammasome”^23^. Here, proteolytically active CASP1 cleaves, and thereby activates pro-inflammatory cytokines IL-1β and IL-18 as well as the pore-forming protein GSDMD. GSDMD pores permeabilize the plasma membrane to facilitate secretion of IL-1β and IL-18^24,25^. DPP8 and DPP9 are required to maintain CARD8 and NLRP1 sensors in an inactive conformation^19–22^. However, DPP8 and DPP9 also function in inflammasome-independent contexts^26^. No studies have mechanistically determined how DPP9 contributes to severe COVID-19 or IPF to date. The critical role of DPP9 in human lung disease, coupled with its established function in regulating the CARD8 and NLRP1 inflammasomes, positions DPP9 as a prime target for investigation in CIP.

In this study, we investigate the contribution of DPP9 to CIP by leveraging large clinico-genomic cohorts, analyses of human immune cells and a genetically engineered mouse model. We performed a stepwise, hypothesis-driven study beginning with a prespecified genetic signal at the *DPP9* locus nominated by prior genome-wide associations with IPF and severe COVID-19. We demonstrate that *DPP9* is a novel CIP susceptibility gene that functions in human monocytes by restricting CARD8-dependent IL-18 secretion. Concordantly, IL-18 was elevated in the plasma of patients receiving ICI-therapy that subsequently developed CIP, suggesting that circulating levels of IL-18 may serve as a novel biomarker that predicts this toxicity. In mice, myeloid-specific deletion of Dpp*8* and *Dpp9* causes spontaneous IL-18-dependent lung inflammation that resembles the immunophenotypic and histopathological features of CIP. This phenotype is fully abrogated by co-deletion of *Il18*. Collectively, the convergence of human genetic, cellular, and *in vivo* murine evidence supports a causal model in which germline-encoded decreased DPP9 function in monocytes lowers the threshold of activation for the CARD8–CASP1–IL-18 axis, predisposing individuals to develop CIP during ICI therapy.

## Results

### DPP9 variants associate with increased risk of CIP

To assess the prevalence of *DPP9* GWAS risk SNPs (rs12610495, rs2109069 and rs2277732) associated with IPF and severe COVID-19, we turned to the gnomAD v4.1 dataset^27^. Analyzing 152,430 alleles, we found that these 3 risk-associated SNPs are common with allele frequencies ranging from 0.24-0.27 when averaged across ancestries (**Fig. 1a, Extended Data Fig. 1a,b**).

**Figure 1.**
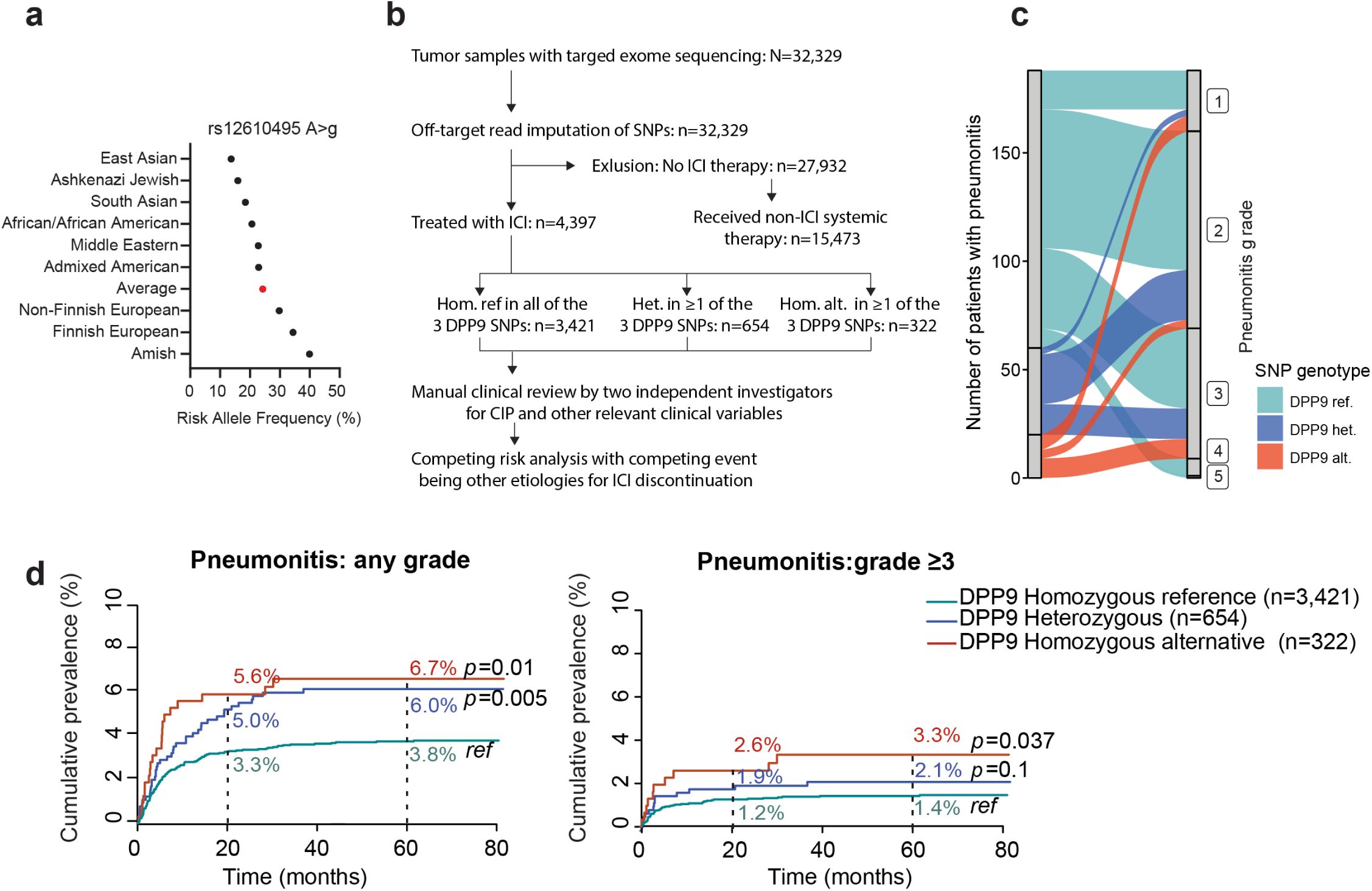
Elevated risk of checkpoint inhibitor pneumonitis (CIP) among carriers of *DPP9* risk alleles. a) Risk allele frequencies of *DPP9* single nucleotide polymorphism (SNP) rs12610495 across nine ancestral populations, as reported in the Genome Aggregation Database version 4.1 (gnomAD v4.1). b) CONSORT diagram of the Dana-Farber Cancer Institute (DFCI) cohort. The three DPP9 SNPs represent previously validated DPP9 risk variants from lung disease GWAS and are analyzed jointly because they tag a common high-linkage disequilibrium haplotype rather than independent association signalsc) Alluvial plot of DFCI cohort demonstrating the number of patients with each grade of pneumonitis based on genotype. d) Cumulative incidence curves for CIP following initiation of immune checkpoint inhibitor (ICI) therapy, estimated using the nonparametric Aalen–Johansen method. Patients in the DFCI cohort are stratified by SNP dosage. CIP is the primary event of interest, with discontinuation of ICI therapy for reasons unrelated to CIP serving as the competing event. The left panel depicts the incidence of CIP of any grade, while the right panel focuses on CIP of Grade 3 or higher. Red curves indicate patients homozygous for at least one of the three *DPP9* SNPs; blue curves represent heterozygous carriers; green curves depict patients lacking these risk alleles. ref: reference

To investigate the association of *DPP9* risk alleles with CIP, we turned to the Dana-Farber Cancer Institute (DFCI) dataset of 4,397 patients treated with ICIs (**Fig. 1b, Supplementary Table 1-2**). Within this cohort, CIP was observed in 4.29% of patients (n=189). Since this cohort size was insufficiently powered to analyze by GWAS, we adopted a hypothesis-driven strategy and tested for an association between CIP and *DPP9* risk SNPs. Specifically, only the three DPP9 SNPs (rs12610495, rs2109069 and rs2277732) previously identified in genome-wide significant associations with IPF and severe COVID-19 were analyzed. These SNPs were selected and the analytic plan finalized before any outcome modeling was performed; no other variants were screened, and no post-hoc filtering based on effect size or significance was applied.

Sex, age at the time of ICI treatment, continental ancestry, and cancer type were adjusted for in the cause-specific Cox proportional hazards model, and Fine-Gray subdistribution hazard model (sHR). CIP risk was assessed as the primary outcome. Discontinuation of immunotherapy due to non-pneumonitis events, along with other clinical factors prompting treatment cessation, were treated as competing risks (see Methods). Direct modeling of competing risks is crucial for accurate incidence computation, as it mitigates potential survivor bias, where individuals with longer survival times may develop CIP by chance^13^.

On multivariable analysis, the sHR model revealed a significantly higher risk of any grade CIP among patients carrying either homozygous (n=322, HR=1.67 [95% CI: 1.03-2.72], *p*=0.01) or heterozygous variants (n=654,HR=1.63 [95% CI:1.13-2.33], *p*=0.008) of *DPP9* SNPs compared to the reference allele (n=3,421, **Fig. 1c-d, Supplementary Table 4**). Moreover, there was a significantly higher risk of ≥grade 3 CIP among patients carrying homozygous (n=322, HR=2.1 [95% CI: 1.04-4.36, *p*=0.037) *DPP9* SNPs compared to the reference allele (n=3,421, **Fig. 1c-d, Supplementary Table 5**).

In an orthogonal approach using SNP dosage as a continuous score, a cause-specific hazard model confirmed a significant association between each DPP9 SNP and ≥grade 3 CIP (rs12610495: HR=1.91 [95% CI: 1.21-3.02], p=0.0054; rs2109069: HR=1.95 [95% CI: 1.25-3.02], p=0.003; rs2277732: HR=1.99 [95% CI: 1.28-3.1], p=0.0023, **Supplementary Table 6**). The three variants reside in strong linkage disequilibrium (pairwise D′ ≈ 0.99–1.00; r² ≈ 0.88–0.95), defining a single DPP9 risk haplotype, and their near-identical effect sizes support a locus-level association rather than SNP-specific effects. Effect estimates were fully concordant in the European ancestry subset (n=4,168), excluding population stratification as a confounder (**Supplementary Table 7**). Notably, *DPP9* SNPs showed no significant association with ≥grade 3 colitis (HR 1.23–1.25, 95% CI crossing 1, p=0.24–0.25) or hepatitis (HR 1.03–1.08, 95% CI crossing 1, p=0.84–0.92), suggesting the DPP9 risk haplotype specifically increases susceptibility to lung toxicity rather than reflecting a broader tendency toward immune-related inflammation. Together, these data nominate DPP9 as the first genetic risk factor for CIP.

### DPP9 restricts CARD8 activation and IL-18 secretion in human monocytes

To identify the cellular compartment in which DPP9 influences CIP development, we examined which pulmonary cell types co-express *DPP9* and the requisite inflammasome machinery. We re-analyzed a single cell RNA-sequencing (scRNA-seq) dataset derived from the lungs of 28 human donors who had no diagnosed lung disease^28^ and found that *DPP9* was expressed widely across cell populations along with the closely related *DPP8* (**Fig. 2a**). *CARD8* and *NLRP1* exhibited more cell-type restricted expression patterns and were highly expressed in multiple myeloid cell types. Consistent with previous studies^29–32^, *NLRP1* was also detected in bronchiolar epithelial cells, and both *CARD8* and *NLRP1* were expressed in lymphoid cells and vascular endothelium. Other inflammasome components including *PYCARD*, *CASP1*, *IL1B* and *IL18* were most prominently expressed in myeloid lineages while *GSDMD* was widely expressed (**Fig. 2a**). The unique co-expression of essential inflammasome machinery within myeloid cells strongly implicated them as key mediators of DPP9-dependent inflammasome regulation. Importantly, we found that rs12610495 “g” is also associated with lower *DPP9* transcript abundance in human monocyte-derived dendritic cells (DCs, **Fig. 2b**, data reanalyzed from^33^). This risk allele may therefore reduce *DPP9* expression to cause dysregulated inflammasome responses in human myeloid cells. To further delineate this relationship, we examined protein-abundance of DPP9, CARD8 and NLRP1 in human primary monocytes, macrophages and DCs. As expected, DPP9, CARD8 and NLRP1 protein was detectable in each cell type (**Fig. 2c**). In monocytes, CARD8 protein was exclusively present as a lower molecular weight protein product corresponding to isoform 1, while isoforms 4 and 5 predominated in macrophages and DCs. These isoforms represent differences in the regulatory amino-terminus of CARD8^34^, but the impact of each isoform on CARD8 function remains incompletely understood.

**Figure 2.**
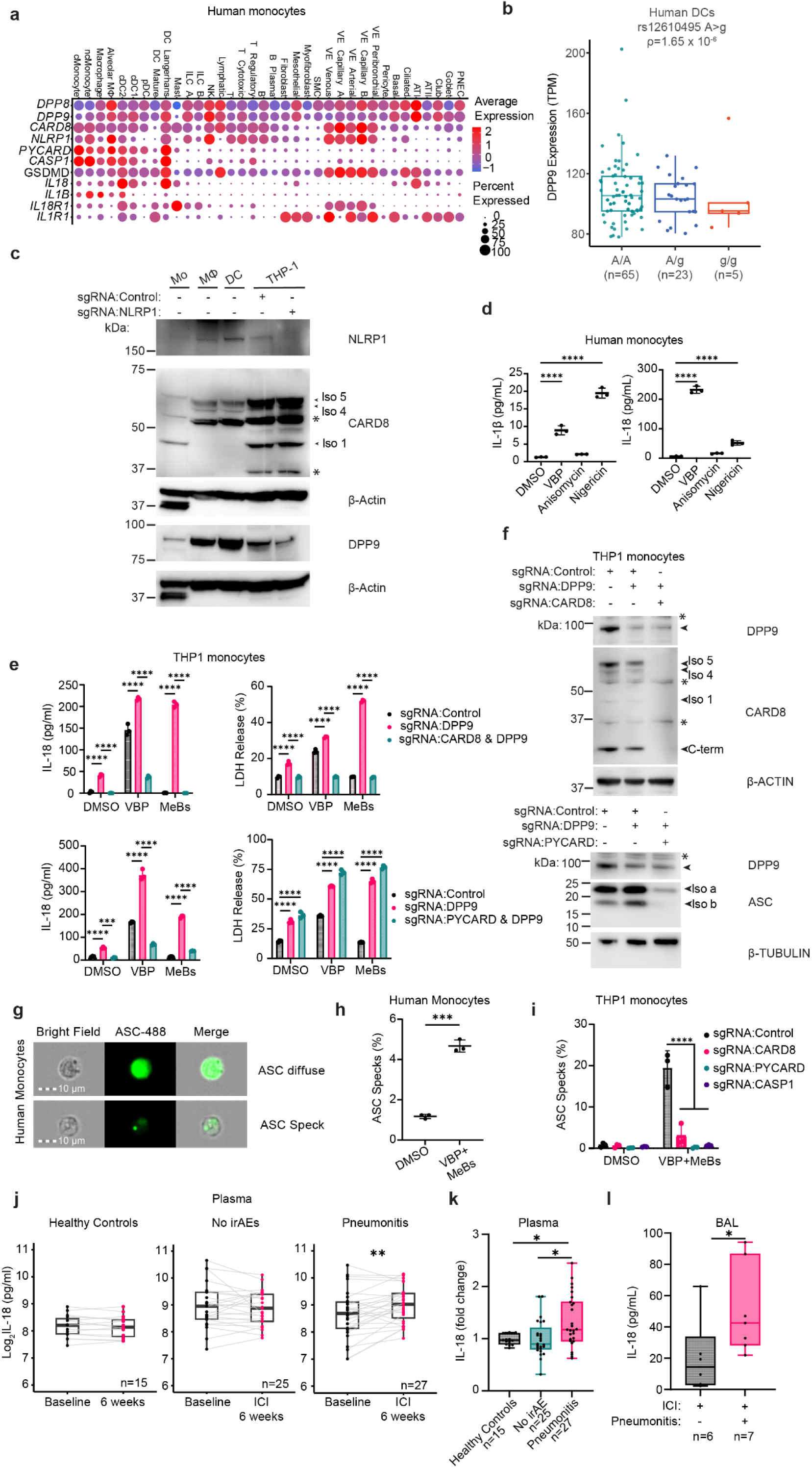
Human DPP9 restricts CARD8 activation and IL-18 secretion in monocytes. a) Dot plot of human lung single cell RNA sequencing data from^28^ indicating mRNA expression levels of inflammasome components. b) DC eQTL data (reanalyzed from^33^) shows rs12610495 “g” allele is associated with reduced *DPP9* expression. Box plot shows median, 25^th^ to 75^th^ percentiles, whiskers mark 1.5 x interquartile range. c) Immunoblot of protein lysates from human monocytes (Mo), macrophages (Mϕ) and dendritic cells (DCs). Non-specific (*) or bands of interest (arrowhead) are indicated. Cas9:sgRNAs complexes targeting Control (*TRAC*) or *NLRP1* were electroporated into THP1 monocytes to validate antibody specificity. d) IL-1β cytometric bead array and IL-18 ELISA of supernatants from primary human CD14^+^ monocytes. One-way ANOVA, Tukey’s multiple comparison test. e) IL-18 ELISAs and LDH release of supernatant from monocytic THP1 cells expressing Cas9 were engineered by electroporating sgRNAs targeting Control (*TRAC*), *DPP9* and *CARD8*. Two-way ANOVA, Tukey’s multiple comparison test. f) Immunoblot of THP1 monocyte protein lysates used to confirm knock-out efficiency. Non-specific (*) or bands of interest (arrowhead) are indicated. g) Representative imaging cytometry data of ASC immunostaining (green) in primary human monocytes. Diffuse and “speck” distribution represent inactive and activated/oligomerized state. h-i) Quantification of imaging cytometry data showing percentage of ASC specks in given treatments in human monocytes and THP1 monocytes, respectively. h) two-tailed unpaired T-test. i) Monocytic THP1 cells expressing Cas9 were engineered by electroporating sgRNAs targeting Control (*TRAC*), *CARD8, PYCARD* or *CASP1,* Two-way ANOVA, Sidak’s multiple comparison test. j-k) IL-18 ELISA data from plasma of indicated patient populations. Levels at baseline and six weeks of ICI therapy are shown, healthy controls received no treatments. j) Box plot shows median, 25^th^ to 75^th^ percentiles, whiskers mark minimum to maximum with grey lines connecting paired patient samples, two-tailed paired t-test and k) box plot shows median, 25^th^ to 75^th^ percentiles, whiskers mark minimum to maximum and each dot represents the fold change for a unique patient, one-way ANOVA, Tukey’s multiple comparison test. l) IL-18 ELISA data from bronchioalveolar lavage fluid (BAL) in patients receiving ICI therapy with (+) and without (-) CIP. Data is visualized as box plot with median, 25^th^ to 75^th^ percentiles, whiskers mark minimum to maximum, each dot is data from a different patient, two-tailed unpaired Mann-Whitney test. Cell culture experiments are representative of at least three independent experiments. Mean is shown with standard deviation of biological replicates corresponding to independently treated wells (n=3). \*\*\*\**p*<0.0001, \*\*\**p*<0.001, \**p*<0.05. Valboro-pro (VBP) 25 µM, 10 µM anisomycin, 10 µM methyl ester bestatin (MeBs) 10 µM nigericin treated 24h. cMonocyte, (classical), ncMonocyte (non-classical), cDC (conventional), pDC (plasmacytoid), VE (vascular endothelial), ATI (alveolar type I), ATII (alveolar type II), PNEC (pulmonary neuroendocrine) see^27^ for further detail. Iso. (isoform), irAE (immune-related adverse events), BAL (bronchioalveolar lavage).

Based on these RNA and protein expression patterns, we hypothesized DPP9 may influence CIP development by restricting CARD8 or NLRP1 activation in human myeloid cells. Treatment of CD14^+^ monocytes with the DPP8/9 inhibitor Valboro-pro (VBP) resulted in robust secretion of the inflammasome regulated cytokine IL-18 and modest secretion of IL-1β (**Fig. 2d**). Interestingly, the CARD8 inflammasome preferentially activates IL-18 in endothelial cells^31^ and has a limited ability to process IL-1β^19,35^. Therefore, we hypothesized that CARD8 mediates VBP-induced IL-18 secretion in human primary monocytes. Treatment of monocytes with anisomycin, a NLRP1-specific activator^36^, did not induce the secretion of inflammasome-regulated cytokines, suggesting that CARD8, but not NLRP1, is functional in human monocytes. In these experiments, the NLRP3 activator nigericin was used as a positive control for cytokine detection (**Fig. 2d**). To confirm that DPP9 restricts CARD8 activation in human monocytes, we applied a genetic approach in human monocytic THP1 cells using CRISPR/Cas9. DPP9-deficient cells exhibited spontaneous cell death and IL-18 secretion that was CARD8-dependent (**Fig. 2e**). We also found that DPP9 was required to prevent CARD8-dependent cell death and IL-18 secretion in response to methyl ester bestatin treatment (MeBs), a known CARD8-activator that induces protein folding stress^37^. In DPP9-deficient THP1 monocytes, MeBs treatment caused cell death and IL-18 secretion in a CASP1-dependent fashion while NLRP1 and NLRP3 were dispensable for these processes (**Extended Data Fig. 2a-c.**) Consistent with previous studies^19,35^, a slight increase in IL-1β secretion was only detectable in DPP9-deficient monocytic THP1 cells treated with MeBs (**Extended Data Fig. 2d**). As expected, VBP treatment caused GSDMD cleavage in a CARD8-dependent fashion in THP1 monocytes (**Extended Data Fig. 2e**). These experiments indicate that DPP9 functions in human monocytes by restricting CARD8-mediated IL-18 secretion.

The limited ability of CARD8 to induce IL-1β secretion is often explained by a model where the CARD8 inflammasome assembles independently of the ASC adaptor protein^38^. Surprisingly, we observed that CARD8-induced IL-18 secretion required ASC (gene name *PYCARD*, **Fig. 2e-f**). However, CARD8-induced cell death occurred independently of ASC (**Fig. 2e-f**), consistent with previous studies^38^. Using imaging cytometry, we confirmed that CARD8 activation led to ASC oligomerization, visualized as a single “speck” in human primary monocytes and monocytic THP1 cells (**Fig. 2g-i, Extended Data Fig. 3**). ASC-speck formation was abrogated in THP1 cells which lacked CARD8 or ASC (**Fig. 2i**). Importantly, CASP1 was also required for ASC oligomerization in response to CARD8 activation (**Fig. 2i**). Therefore, we propose that CASP1 recruits ASC to the CARD8 inflammasome. Notably, previous studies that demonstrated CARD8 does not associate with ASC relied on ectopic expression systems in cells that did not express CASP1^38^.

Surprisingly, VBP treatment did not lead to secretion of inflammasome-regulated cytokines in human primary macrophages or DCs (**Extended Data Fig. 4a,b**) suggesting that monocytes are exceptionally sensitive to DPP9 activity levels compared to other human myeloid cells. Anisomycin treatment triggered IL-1β and IL-18 secretion from human DCs (**Extended Data Fig. 4b**). Using CRISPR/Cas9-mediated gene targeting, we also confirmed that anisomycin-induced cytokine secretion depended on NLRP1 and CASP1 in DCs (**Extended Data Fig. 4c,d**). Collectively, these findings establish DPP9 as a key repressor of CARD8-mediated IL-18 secretion in human monocytes, a mechanism with potential relevance to inflammatory lung conditions.

Having established that DPP9 loss unleashes CARD8-dependent IL-18 secretion in human monocytes, we asked whether this cytokine axis is engaged in patients who develop CIP. Precedent for IL-18 as a driver of pulmonary pathology exists in IPF, where elevated IL-18 has been detected in both serum and bronchoalveolar lavage (BAL) fluid^39^. Consistent with a potential causal role in CIP, plasma IL-18 rose significantly above pre-treatment levels after 6 weeks of ICI therapy in patients who subsequently developed pneumonitis, but not in ICI-treated patients who remained free of immune-related adverse events (**Fig. 2j-k, Supplementary Table 9**). Critically, this elevation preceded the clinical diagnosis of pneumonitis in 85% of patients, raising the possibility that circulating IL-18 could serve as an early predictive biomarker of CIP. Corroborating this systemic signal at the site of disease, IL-18 protein was also significantly elevated in BAL fluid from ICI-treated patients with CIP relative to those without pneumonitis (**Fig. 2l**, **Supplementary Table 10**). These clinical findings position IL-18 as a predictive biomarker and potential mediator of CIP.

### Murine DPP8 and DPP9 prevent NLRP1-driven pulmonary inflammation

Pathogen activators of NLRP1 and CARD8 are species-specific for humans or rodents; however, regulation through DPP9 is a evolutionarily conserved between humans, rodents and zebrafish^19,20,22,40,41^. CARD8 has been described as a “NLRP1-like” inflammasome sensor with similar mechanisms of regulation^42^. Since mice lack CARD8, we hypothesized that murine NLRP1 may serve a function analogous to human CARD8 when regulation by DPP9 is compromised. Mice possess three tandem paralogs of *Nlrp1*(a-c), of which, NLRP1a and NLRP1b are activated by DPP9 inhibition^43,44^. To determine if NLRP1a/b respond to DPP9 inhibition in mouse monocytes, we utilized mice engineered with a germline knock-out of *Nlrp1b* generated in a haplotype where *Nlrp1a* and *Nlrp1c* are not functional^44,45^. Treatment of primary mouse monocytes with the DPP9 inhibitor VBP led to NLRP1-dependent secretion of inflammasome regulated cytokines IL-1β and IL-18 as well as GSDMD cleavage (**Fig. 3a,b**). Thus, murine NLRP1 is activated in response to DPP9 deficiency in monocytes, serving a function that is analogous to CARD8 in human monocytes.

**Figure 3.**
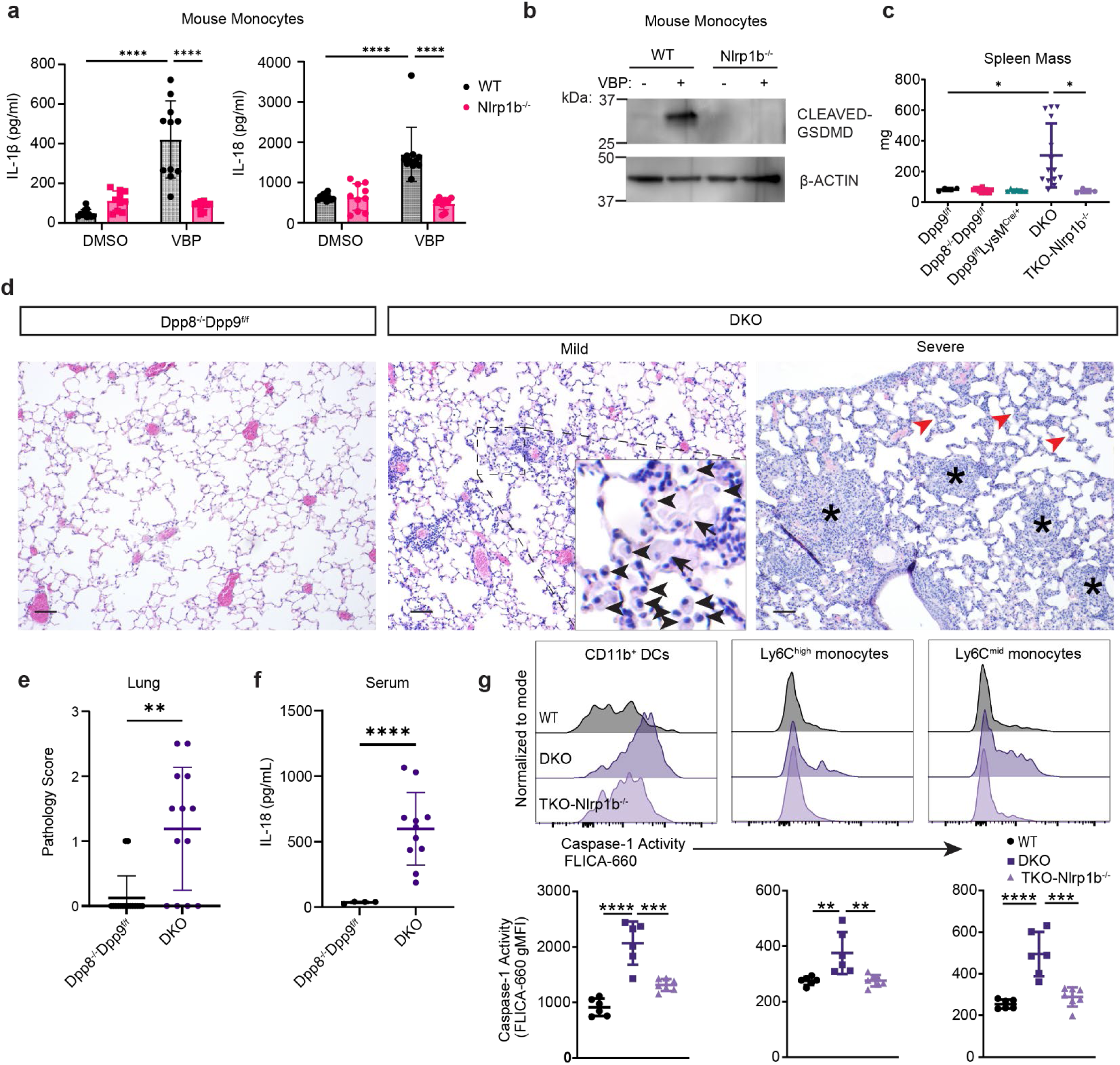
DPP8 and DPP9 function in myeloid cells to prevent NLRP1 driven CIP-like lung inflammation. a) ELISA and b) immunoblot data of primary mouse monocytes with indicated genotypes and treatments. Two-way ANOVA, Tukey’s multiple comparison test. Valboro-pro (VBP) 25 µM treated 24 h (n=10-11). c) Spleen weights from indicated genotypes (n=4-14). One-way ANOVA, Tukey’s multiple comparison test. d) Hematoxylin and Eosin (H&E) staining of the lungs in indicated genotypes (n=13-16). Mild and severe DKO phenotypes are shown. Alveolar macrophages and foamy cells are highlighted by black arrowheads and arrows, respectively. Granuloma-like structures and alveolar thickening are marked by asterisks and red arrowheads, respectively. Scale bar is 100 µm. e) Lung pathology scores analyzed using two-tailed unpaired T-test (n=13-16). f) IL-18 ELISA data from serum of mice with indicated genotypes analyzed using two-tailed unpaired T-test (n=4-11). g) Flow cytometry quantification of FLICA-660 CASP1 reporter probe in indicated genotypes and cell populations (n=6). Representative histograms and quantification. One-way ANOVA, Tukey’s multiple comparison test. Mean is shown with standard deviation. For *in vivo* experiments, each dot represents values from a different mouse. \*\*\*\**p*<0.0001, \*\*\**p*<0.001, \*\**p*<0.01, \**p*<0.05. All results are representative of at least three independent experiments. DKO, double knock-out (*Dpp8^-/-^Dpp9^f/f^LysM^Cre/+^*) TKO, triple knock-out.

To determine how DPP9 functions in myeloid cells to regulate pulmonary inflammation, we generated mice that harbor a germline mutation in *Dpp8* or a floxed-allele of *Dpp9* (*Dpp9^f^*) which was crossed to the LysM-Cre^46^. The germline mutation in *Dpp8* was introduced to address the potential functional redundancy^20,35^ between DPP8 and DPP9 as has been observed in the regulation of human NLRP1^20^. Strikingly, we observed splenomegaly and lymphadenopathy specifically in mice that were *Dpp8 and Dpp9* double knock-out (DKO), but not either single mutant (**Fig. 3c, Extended Data Figure 5a**). This phenotype did not manifest in triple knock-out mice deficient in *Dpp8*, *Dpp9*, and *Nlrp1b* (TKO-*Nlrp1b^-/-^*) (**Fig. 3c**), indicating these spontaneous phenotypes require NLRP1. DKO mice exhibited gross inflammation that primarily involved the lung and liver (**Fig. 3d**, **Extended Data Fig. 5a**). The extent of inflammation ranged from minimal to severe, with rare instances of spontaneous fibrosis in the lung and liver (**Fig. 3d-e**, **Extended Data Fig. 5a-b**). Pulmonary inflammation in DKO mice resembled histopathologic features previously described in patients with CIP^47^. Of note, mildly affected DKO mice possessed alveolar regions with a localized high density of macrophages, including foamy cells (**Fig. 3d**). More severe cases of pulmonary inflammation presented with poorly-defined granulomas and alveolar thickening. This pattern of pulmonary inflammation is also consistent with an immune-mediated non-infectious etiology like CIP. Serum IL-18 levels were also elevated in DKO mice, although IL-1β abundance was similar in both genotypes (**Fig. 3f, Extended Data Fig 5d**). Despite spontaneous inflammation, DKO mice-maintained survival rates comparable to littermate controls (**Extended Data Fig. 5e**). These data suggest that DPP8/9 function in myeloid cells by preventing spontaneous CIP-like inflammation, especially in the lung.

To assess inflammasome activity in pulmonary myeloid cells, we utilized the FLICA CASP1 reporter. Increased FLICA signal was observed in Ly6C^high^ and Ly6C^mid^ monocytes as well as CD11b^+^ DCs from DKO mice (**Fig. 3g**). In each of these cell types, levels of CASP1 activity were reduced to the level of control mice in TKO-*Nlrp1b^-/-^*mice, showing that increased CASP1 activity was driven by NLRP1. These results indicate that in the absence of rodent homologs for CARD8, murine NLRP1 compensates by fulfilling the roles of human CARD8 in monocytes and NLRP1 in DCs. Further, constitutive NLRP1-mediated inflammasome activation in mouse myeloid cells leads to multiorgan inflammation influencing the lung, liver, spleen and lymph nodes. These findings also establish a causal relationship between loss of DPP8/9-mediated inflammasome suppression and pulmonary inflammation that resembles CIP histologically.

### The murine NLRP1-CASP1-IL-18 pathway promotes a pulmonary type I T cell response

Our next objective was to investigate whether DPP8/9 deficiency and NLRP1 signaling drives a pulmonary immune program like that observed in patients with CIP. We profiled lungs of WT, DKO and severe DKO mice by scRNA-seq, and analyzed 14,482 high quality cells from these three groups (WT: 4,995, DKO: 1,691, DKO-severe: 7,796, **Extended Data Fig. 6-7)**. DKO, and to a greater extent severe DKO, mice exhibited distinct clusters of CD4^+^ T cells expressing *Cxcr3*, and *Ifng* as well as CD8^+^ T cell clusters characterized by high expression of *Ifng*, *Ccl4*, *Ccl5* and *Gzmb*, indicative of an active type I immune response (**Fig. 4a,b**, **Extended Data Fig. 8a,b**). Flow cytometry analysis corroborated that DKO mice displayed an increased presence of antigen-experienced CD44^+^ CD4^+^ and CD8^+^ T cells in the lung (**Extended Data Fig. 8c**). Further, a higher percentage of DKO-derived CD4^+^ and CD8^+^ T cells produced IFNγ upon *ex vivo* stimulation (**Fig. 4c)**, a hallmark previously observed in patients with CIP^9,10^. T cell composition and cytokine production were comparable in control, *Dpp8* and *Dpp9* single mutant mice indicating these negative regulators function redundantly in mice (**Fig. 4c, Extended Data Fig. 8c**). Analysis of multiple TKO mouse lines indicated that CD4^+^ and CD8^+^ T composition and IFNγ production were contingent on the presence of NLRP1, CASP1 and IL-18 (**Fig. 4c, Extended Data Fig. 8c**), while IL-1 is not required as the IL-1β receptor IL1R proved dispensable for these observed T cell phenotypes. Finally, analysis of serum cytokines by multiplexed ELISA identified IFNγ as the most upregulated factor within a panel of 27 secreted proteins indicating T cells were activated *in vivo* (**Fig. 4d**).

**Figure 4.**
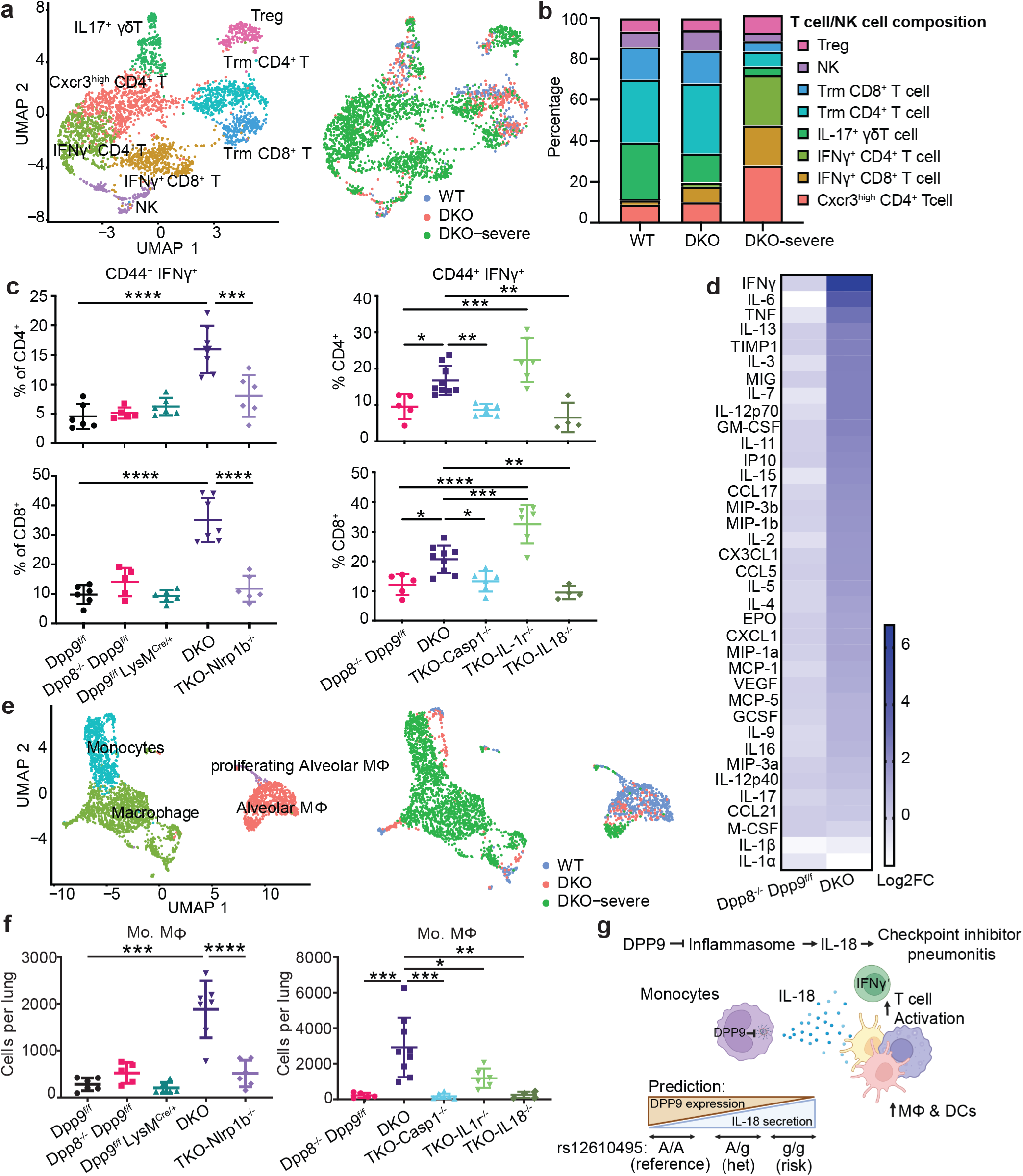
DPP8 and DPP9 prevent spontaneous type I immunity driven by the NLRP1 inflammasome and IL-18. a) scRNA-seq data from WT, DKO and DKO-severe shown as UMAP of T/NK cell subclusters and indicated genotypes. b) Stack plot showing frequencies of each T/NK subcluster by genotype. c) Flow cytometry quantification of IFNγ production from *ex vivo* stimulated CD4^+^ and CD8^+^ T cells from indicated genotypes (n=4-9). d) Multiplex ELISA from serum samples of indicated genotypes (n=6-8). e) scRNA-seq data from WT, DKO and DKO-severe shown as UMAP of myeloid cell subclusters and indicated genotypes. f) Flow cytometry quantification of monocyte derived macrophages (Mo. Mϕ) from indicated genotypes (n=4-9). g) Proposed model: DPP9 suppresses inflammasome activation and IL-18 secretion from monocytes. IL-18 promotes a CIP-like type I immune response characterized by spontaneous IFNγ secretion and increased abundance of inflammatory macrophage (Mϕ) and DC infiltrates (biorender). For flow cytometry data, mean is shown with standard deviation of biological replicates. Each dot represents values from a different mouse. Flow cytometry data was analyzed with a One-way ANOVA, Tukey’s multiple comparison test. \*\*\*\**p*<0.0001, \*\*\**p*<0.001, \*\**p*<0.01, \**p*<0.05. All flow cytometry results are representative of at least two independent experiments. DKO, double knock-out (*Dpp8^-/-^Dpp9^f/f^LysM^Cre/+^*) TKO, triple knock-out.

scRNA-seq data also indicated DKO mice had altered composition of myeloid cells characterized by an increase in monocyte-derived macrophages (**Fig. 4e**). Flow cytometry further confirmed that the lungs of DKO mice harbored a higher abundance of monocyte-derived macrophages, alongside type I and type II DCs, identified by the markers XCR1 and SIRPα, respectively (**Fig. 4f, Extended Data Fig. 11).** This mirrors the increase in pro-inflammatory macrophage and DC populations documented in BAL fluid from patients with CIP^9,10^. Immunophenotyping of TKO mouse lines indicated that the observed increase in macrophages and DCs was dependent on NLRP1, CASP1 and IL-18 (**Fig.4f, Extended Data Fig. 11b**).

These immunophenotyping results demonstrate that murine myeloid-specific DPP8/9 deficiency causes a CIP-like immune response that is mediated by the inflammasome and IL-18. Together with our clinico-genomic and human immune cell studies, our findings suggest that *DPP9* SNPs increase an individual’s CIP risk by impairing repression inflammasome-mediated IL-18 secretion in human monocytes, thereby promoting an IFNγ predominant T cell phenotype associated with a high risk of mortality (**Fig. 4g**).

## Discussion

Our study offers pivotal insights into CIP, an immune related adverse event that leads to an estimated 20,000 deaths in the United States annual^48–50^. By identifying a correlation between *DPP9* disease-associated SNPs and increased CIP risk, we have uncovered a key genetic factor that could be crucial for predicting and managing this life-threatening condition. We show that *DPP9* risk alleles are associated with lower gene expression in myeloid cells and propose that this dysregulation may cause CIP pathogenesis through excessive IL-18 secretion. However, the exact role of these SNPs in DPP9-mediated immune regulation remains to be fully understood^14,17^. Our results position DPP9 as a critical biomarker for stratifying patients undergoing ICI treatment, paving the way for more personalized ICI therapies designed to reduce the risk of severe immune-related adverse events.

We demonstrate that while significant details vary between mice and humans, at least three important features of this pathway are conserved between species. Specifically, 1) human DPP9 and murine DPP8/9 are essential for restricting myeloid-derived IL-18 secretion and 2) patients with CIP and DKO mice exhibit pulmonary inflammation characterized by more IFNγ-producing T cells, inflammatory macrophages and DCs^9,10^. 3) We also show that IL-18 secretion was higher in patients with CIP prior to diagnosis and that in DKO mice, IL-18 was essential for the observed spontaneous type I T cell response. This highlights that IL-18 blockade may be a promising therapeutic strategy for CIP, where morbidity and mortality are high, and treatments are limited to broadly acting immunosuppressants.

Beyond the lungs, our study demonstrates that murine DPP8/9 play a crucial role in preventing inflammasome-driven inflammation in the liver, spleen, and lymph nodes, even though these extra-pulmonary manifestations are not commonly observed in patients with CIP^51^. However, similar systemic inflammation has been documented in an individual with a *de novo DPP9* mutation^52^. Notably, this patient exhibited a ten-thousand-fold upregulation of IL-18 and was diagnosed with hemophagocytic lymphohistiocytosis-like disease that is typically caused by excessive IFNγ production^52,53^. This highlights a broader role of DPP9 in maintaining immune homeostasis across multiple organ systems.

Our human studies have limitations. Because the DFCI cohort represents a retrospective clinico-genomic analysis leveraging tumor sequencing performed as part of routine care under broad research consent, we lack granular data on active outreach (e.g. patients approached, refusals, or failed sequencing attempts), which precludes a complete reconstruction of the participation flow beyond available sequencing and treatment records. Accordingly, the analytic cohort is inherently conditioned on tissue availability and completion of clinical molecular profiling at diagnosis. As such, the DFCI cohort may be enriched for patients with adequate diagnostic tissue and sufficient clinical stability to undergo biopsy and sequencing, an important potential source of selection bias that could influence the observed associations. In addition, the DFCI cohort is predominantly composed of individuals who self-identified as non-Hispanic White, reflecting the institutional catchment area and prior DFCI-based genomic studies. Although adjustment for race/ethnicity and restriction to European ancestry yielded effect estimates for *DPP9* risk alleles that were highly concordant with the primary analysis, this demographic imbalance limits the generalizability of effect size estimates to more diverse populations and underscores the need for validation in ancestrally heterogeneous cohorts. Finally, although the three *DPP9* variants examined lie within a single high-linkage-disequilibrium haplotype and represent a robust locus-level association with CIP risk, the available event count limits statistical fine-mapping, precluding resolution of a uniquely causal variant within the locus and motivating future studies combining larger cohorts with functional dissection of regulatory elements.

In conclusion, our study positions DPP9 as a critical regulator of immune dynamics, directly linking genetic predisposition to the development of CIP. By elucidating the intricate role of DPP9 in suppressing inflammasome-mediated secretion of IL-18, we have identified IL-18 as a promising therapeutic target with the potential to enhance the safety of cancer immunotherapy. These findings constitute a major advance in elucidating the molecular drivers of CIP, laying the foundation for personalized treatment strategies that could markedly enhance the management of immune-related adverse events in cancer care. Given the current dependence on broadly immunosuppressive corticosteroids as the primary treatment for CIP, IL-18 blockade represents a mechanistically rational and potentially more precise alternative. More broadly, this work illustrates how hypothesis-driven integration of clinico-genomic and experimental evidence can transform a genetic association into an actionable biological insight, an approach that may serve as a model for dissecting the genetic architecture of other challenging immune-related adverse events.

## Methods

### Cohort definition, consent and genotyping

This study was conducted in strict accordance with all applicable ethical guidelines. Analyses were performed on the Dana-Farber Cancer Institute (DFCI) cohort, which included comprehensive genotyping and clinical data.

### DFCI cohort

Between 2012 and 2024, a cohort of 4,397 patients at DFCI received treatment with immune checkpoint inhibitors (ICIs) (**Fig. 1b**), representing 14 cancer types with at least 100 patients per type, as well as a combined category for rarer tumors. Approximately 92% of these patients were treated with PD-1/PD-L1 inhibitors, while around 8% received combination immunotherapy, defined by the concurrent administration of CTLA-4 and PD-1/PD-L1 inhibitors (**Supplementary Tables 1-3**). Tumor samples were collected through biopsy or surgical resection and analyzed using the OncoPanel sequencing platform, which targets 275–447 cancer-relevant genes. Germline single nucleotide polymorphisms (SNPs) were inferred from ultra-low-coverage off-target reads, with imputation accuracy verified against a partially overlapping cohort of directly genotyped individuals. To provide a normative comparison, a pan-cancer control group comprising 27,932 individuals treated with non-ICI therapies at DFCI was sequenced and processed through the same analytical pipeline. Collected clinical data included demographic information, relevant oncologic history, the specific ICI agent administered, and treatment duration. Checkpoint inhibitor pneumonitis (CIP) was diagnosed based on evaluations by the treating oncologist or pulmonologist, with severity classified according to the Common Terminology Criteria for Adverse Events (CTCAE) v5.0. Manual chart reviews were conducted for all 4,397 patients by two independent investigators blinded to SNP status (A.H.N., E.B.F.). The DFCI samples were collected and sequenced under IRB-approved protocols 11–104 and 17–000 from the Dana-Farber/Partners Cancer Care Office for the Protection of Research Subjects, with written informed consent obtained from all participants. Secondary analyses of previously collected data were conducted with the approval of the Dana-Farber IRB under protocols 19–033 and 19–025, both of which included a waiver of Health Insurance Portability and Accountability Act (HIPAA) authorization.

### Genomic Data Acquisition, Imputation, and Quality Control in the DFCI Cohort

The DFCI cohort was sequenced as part of the PROFILE project, a prospective clinical sequencing initiative for consented patients undergoing routine treatment at DFCI and affiliated hospitals. Genomic variation was assessed using a custom hybrid capture sequencing platform (OncoPanel), with each sample sequenced on one of three panel versions targeting the exons of 275, 300, or 447 genes, respectively^54^. Analysis was conducted on samples meeting a minimum coverage of 30X for 80% of targeted regions. Somatic variations, including single-nucleotide variants, insertions/deletions, and copy number variations, were identified using the Profile clinical bioinformatics pipeline and reviewed by a pathologist at Brigham & Women’s Hospital following technical validation, as previously described^54^. Off-target and on-target reads from the sequenced BAM files were imputed using the STITCH software v1.6.6^13,55^. Imputed variants were restricted to those with a minor allele frequency (MAF) > 1% and an imputation INFO score >0.4. Imputation accuracy was benchmarked against a partially overlapping cohort of 833 individuals (126 of whom were treated with ICIs) who had both OncoPanel tumor sequencing and direct germline SNP array genotyping (Illumina MEGA). Pearson correlation coefficients were calculated for each SNP between the tumor-imputed and germline-genotyped data, yielding a mean correlation of 0.86 after variant quality control, with high uniformity across the genome. A detailed analysis of variant imputation accuracy has been described separately, and the imputation workflow is publicly available^13^.

### Selection of *DPP9* variants for analysis

We used a hypothesis-driven, locus-focused strategy to nominate DPP9 variants, rather than data-driven selection within the Dana-Farber cohort. Specifically, rs12610495, rs2109069, and rs2277732 were chosen a priori based on reproducible genome-wide significant associations with IPF and severe COVID-19 in independent studies^14,17,18^. Variants were further required to (i) map within the DPP9 locus, (ii) have sufficient minor allele frequency to support time-to-event modeling, and (iii) meet imputation quality thresholds in the DFCI pipeline (INFO > 0.4; Methods). Because these SNPs reside in strong linkage disequilibrium, we treated them as correlated markers of a shared DPP9 risk haplotype and interpreted results at both the locus level rather than as independent tests. Variant selection was finalized before outcome modeling, and no post hoc filtering was performed based on association statistics in the DFCI cohort. Because the three *DPP9* SNPs are in strong linkage disequilibrium, as assessed using LDlink^56^, and were prespecified as correlated markers of a single locus-level hypothesis, we did not apply multiple-testing correction for independent tests; inference was therefore made at the locus level.

### Patient plasma analysis

Plasma samples were obtained from 67 patients at UT Southwestern Medical Center as previously described^57^, comprising 15 healthy controls with no ICI exposure, 25 patients receiving immune checkpoint inhibitor (ICI) therapy who developed no immune-related adverse events (no irAE), and 27 patients who developed checkpoint inhibitor pneumonitis. Samples were collected and analyzed under protocol STU 082015-053 approved by UT Southwestern Institutional Review Board, with written informed consent obtained from all participants. Samples were collected at baseline (pre-ICI) and 6 weeks post-ICI initiation (**Supplemental Table 9**). Plasma IL-18 concentrations were measured using the Human Total IL-18 DuoSet ELISA Kit (Catalog #DY318-05; Bio-Techne, Minneapolis, MN) per the manufacturer’s instructions, with all samples run in duplicate.

Statistical analysis: Intra-assay precision was assessed by coefficient of variation (CV with only CV < 15% accepted; most CVs ranged from 0.5–2.5% and all standard curves met a minimum R² of 0.997. Prior to analysis, IL-18 concentrations were log₂-transformed, which conformed to a normal distribution. Parametric tests were used throughout data analysis. Within-group changes from baseline to 6 weeks were assessed by paired *t*-test. Fold change was calculated for between-group differences and assessed by one-way ANOVA with Tukey’s multiple comparison test. Data was visualized as box plots. All statistical analyses were performed in GraphPad Prism 10.6.1.

### Patient Bronchioalveolar lavage (BAL) fluid analysis

BAL fluid IL-18 concentrations were measured in an independent cohort of 13 patients who underwent bronchoscopy at Johns Hopkins, as previously described^58^. Patients were categorized into two groups: 6 patients treated with ICI who did not develop pneumonitis (ICI+/CIP−), and 7 patients who developed checkpoint inhibitor pneumonitis (ICI+/CIP+, **Supplementary Table S10**). Samples were collected and analyzed under protocol IRB00186276 approved by Johns Hopkins Hospital Institutional Review Board, with written informed consent obtained from all participants. IL-18 concentrations in BAL fluid were measured using the Human Total IL-18 DuoSet ELISA Kit (Catalog #DY318-05; Bio-Techne, Minneapolis, MN) per the manufacturer’s instructions. IL-18 concentrations are shown as box plots and analyzed by Mann-Whitney test. All statistical analyses were performed in GraphPad Prism 10.6.1.

### Multistate modeling of competing risks in CIP analysis

Two complementary time-to-event analyses were performed with CIP as the primary outcome. In the cause-specific Cox proportional hazards model, patients who died or discontinued ICI therapy for any reason other than pneumonitis were censored at the time of discontinuation. This approach measures the inherent biological risk of CIP among patients who remain event-free, and the cause-specific hazard ratio was therefore the primary metric of interest given our focus on the mechanistic underpinnings of CIP. To formally account for the competing risk of treatment discontinuation, a Fine-Gray subdistribution hazard model was additionally applied, in which pneumonitis was coded as Event=1, any non-pneumonitis discontinuation (such as death, disease progression, and other clinical factors prompting ICI cessation) as Event=2, and loss to follow-up as Event=0. Direct modeling of competing risks is crucial for accurate incidence computation, as it mitigates potential survivor bias, where individuals with longer survival times may develop CIP by chance.

### Statistical analysis

The association of *DPP9* SNPs with time to CIP was evaluated across the DFCI cohort. CIP probabilities and cumulative incidences were calculated using the Aalen–Johansen estimator, a nonparametric approach that accounts for competing risks^59^. Given the competing risk of death during treatment, cause-specific hazard ratios (HRs) were computed for each SNP using a mixed-effects model, effectively censoring on death or loss to follow-up. Each cohort’s analysis included covariates for age at ICI initiation, sex, cancer type (lung, melanoma, genitourinary, gastrointestinal, breast, sarcoma/other), and type of ICI (PD-1/PD-L1 or CTLA4 monotherapy or combination). CIP probabilities and cumulative incidences were quantified using the Aalen–Johansen estimator, consistent with the competing risks framework^59^.

### eQTL analysis

Bulk RNA-sequencing data was reanalyzed from 99 monocyte-derived dendritic cells^33^. Monocytes were cultured for 7 d in RPMI (Life Technologies) supplemented with 10% FBS, 100 ng/mL GM-CSF (R&D Systems), and 40 ng/mL IL-4 (R&D Systems) to differentiate the monocytes into MoDCs; 4 × 10^4^ MoDCs were seeded in each well of a 96-well plate. RNA from all samples was extracted using the RNeasy 96 kit (Qiagen, catalog no. 74182), according to the manufacturer’s protocols and sequenced to an average depth of 38 million 76-bp paired-end reads using the Illumina TruSeq kit. After aligning reads to the genome, transcriptomes were assembled for each sample individually using StringTie (Pertea et al. 2015) and default parameters. Abundances of annotated transcripts were quantified using Kallisto (Bray et al. 2016). As previously described (Lee et al. 2014), genomic DNA was extracted from 5 mL whole blood (DNeasy blood & tissue kit; Qiagen) and quantified by NanoDrop. Each subject was genotyped using Illumina infinium human OmniExpress exome BeadChip, which includes genome-wide genotype data as well as genotypes for rare variants from 12,000 exomes as well as common coding variants from the whole genome. In total, 951,117 SNPs were genotyped, of which 704,808 SNPs are common variants (minor allele frequency [MAF] > 0.01) and 246,229 are part of the exomes. To accurately evaluate the evidence of association signal at variants that are not directly genotyped, we used BEAGLE (Browning and Browning 2016) software (v3.3.2) to impute the post-QC genotyped markers using reference haplotype panels from The 1000 Genomes Project (The 1000 Genomes Project Consortium PhaseI Integrated Release Version 3) (Siva 2008),Local eQTL mapping was performed using the Matrix eQTL (Shabalin 2012) package using empirically determined number of principal components (PCs) as covariates for each analysis. Experiment-wide empirical P-values were calculated by comparing the nominal P-values with null P-values determined by permuting each isoform/gene 1000 times (Churchill and Doerge 1994). The permutation P-values were not pooled to calculate the empirical P-values (i.e., the minimum possible P-value is 0.001). False-discovery rates were calculated using the qvalue package (https://github.com/StoreyLab/qvalue) as previously described (Storey and Tibshirani 2003).

### Cell culture

Human blood was collected from healthy volunteers in BD vacutainer EDTA tubes (BD#366643) according to the principles of the Declaration of Helsinki and was approved by the Yale New Haven Hospital Institutional Review Board (IRB protocol 1507016144). The study relied solely on deidentified data; therefore, written informed consent was waived. Monocytes were purified by negative selection (Stem Cell #9669). Monocytes were differentiated into macrophages or DCs in RPMI supplemented with 10% fetal bovine serum, 100 units/mL penicillin, 100 µg/mL streptomycin and 2mM L-glutamine along with 50 ng/mL human M-CSF (Biolegend #574804) or the combination of 50 ng/mL human GM-CSF (Peprotech #300-03) and 40 ng/mL human IL-4 (Peprotech #200-04). Cells were cultured for two weeks, and the media was changed three times.

Primary mouse monocytes were purified by negative selection from bone marrow (Biolegend, 480154). Cells stimulated for 24 hours in RPMI supplemented with 10% fetal bovine serum, 100 units/mL penicillin, 100 µg/mL streptomycin and 2mM L-glutamine.

### *In vitro* CRISPR/Cas9

Monocytic THP1 cells expressing Cas9 were obtained from the Sidi Chen lab at Yale University and cultured in RPMI supplemented with 10% fetal bovine serum, 100 units/mL penicillin, 100 µg/mL streptomycin and 2 mM L-glutamine. sgRNAs targeting *CARD8* (Synthego) were transfected into THP1 cells using Lipofectamine CRISPRMAX (ThermoFischer CMAX00001) following the manufacturer’s protocol. Single clones were FACs sorted into a 96-well plate, expanded and validated by immunoblot. Polyclonal cell populations were analyzed for DPP9, CASP1, NLRP3, NLRP1 and ASC and knock-out efficiency was assessed by immunoblot. 2×10^6^ Cas9-expressing THP1 cells were electroporated with an RNP complex of 60 pmol Cas9 and 120 pmol of pooled sgRNAs (Synthego), using Amaxa SG Cell Line protocol (Lonza, V4XC-3032) in the Lonza 4D-nucleofector X unit. For DCs, primary human monocytes were edited using the same RNP complex, but using the P3 primary Cell kit (Lonza, V4XP-3032) as previously described^60^.

TRAC: (Negative control, 5’ to 3’)

sg1 (CUCUCAGCUGGUACACGGCA)

sg2 (GAGAAUCAAAAUCGGUGAAU)

sg3 (ACAAAACUGUGCUAGACAUG)

DPP9: (5’ to 3’)

sg1 (GGAUUAGCCCCAUGAGCGAG)

sg2 (GGUGGAGAUCGAGGACCAGG)

sg3 (CAUGGAUGGCAACUCGGCUC)

CARD8: (5’ to 3’)

sg1 (UAAGAUAGGACACGAGGUAA)

sg2 (GACACGGCAGAGCAAGAAUG)

sg3 (CCGGCAACUCCAAGCCAGGA)

NLRP1: (5’ to 3’)

sg1 (UUUCAGGAGGACUCCCAAGG)

sg2 (UAUGUGAUGCAGCUCCACCC)

sg3 (UCCCAAGUGACUGCUCCAUU)

NLRP3: (5’ to 3’)

sg1 (GAGGACUAUCCUCCCCAGA)

sg2 (AGAGGACUAUCCUCCCCAGA)

sg3 (AAUGAUCGACUUCAAUGGGG)

PYCARD: (5’ to 3’)

sg1 (GUUCUCCAGCGCAUCCAGGA)

sg2 (UCAGGUUCUCCAGCGCAUCC)

sg3 (CGACGCCAUCCUGGAUGCGC)

CASP1: (5’ to 3’)

sg1 (UUUAUCCGUUCCAUGGGUGA)

sg2 (CUAAACAGACAAGGUCCUGA)

Chemical Reagents
|  |  |  |  |
| --- | --- | --- | --- |
| LPS | Enzo | ALX-581-012-L002 | 100 ng/mL |
| Talabostat mesylate (VBP) | Medkoo | 202730 | 25 μM |
| Anisomycin | Tocris | 1290 | 10 μM |
| Methyl ester bestatin (MeBs) | Santa Cruz | SC-221350 | 10 μM |
| Nigericin | Cayman | 11437 | 10 μM |
| VX-765 | APExBIO | A8238 | 0.1mg/mL |

### Imaging cytometry

1 x 10^6^ monocytes or THP1 cells were incubated with 0.100 mg/mL of CASPASE-1 inhibitor VX-765 and treated as indicated. VX-765 increases detection of ASC specks by preventing cell death^61,62^. Cells were fixed and permeabilized according to BD cytoplasmic kit (BD 555028), stained with anti-ASC antibody (Cell signaling 78000, 1:100) over night at 4°C then incubated in anti-rabbit AF488 (Invitrogen A21206, 1:1000) secondary antibody for at least 1 hour at room temperature in the dark. Stained cells were analyzed on Amnis Imagestream-X Mark II and quantitated using IDEAS software.

### Immunoblot

Cells were lysed with in RIPA buffer (Millipore 20-188) containing protease inhibitor cocktail (Roche 04-693-159-001). Protein samples were prepared with NuPAGE LDS sample buffer (NP0007) and reducing agent (NP0009) and boiled for 5 min. Nupage 4-12% Bis-Tris gels were used for SDS PAGE electrophoresis with MOPS running buffer (NP0001) and transferred to PVDF membrane (Biorad 1620177) with NuPAGE transfer buffer (NP00061) supplemented with 10% Methanol at 80V for 3 hours. Protein-containing membranes were blocked briefly in 5% milk in PBST (containing 0.1% Tween) and primary antibodies were incubated overnight at 4°C. Membranes were washed three times with PBST then incubated with secondary antibodies for 1 hour at room temperature. For blots that detected NLRP1 and cleaved-GSDMD, antibodies were incubated in “Can Get Signal” (Fisher, NC1518201), otherwise antibodies were 5% milk in PBST was used as an antibody incubation buffer. Blots were developed using Femto ultra-sensitive enhanced chemiluminescent HRP substrate (Thermo#34094) and ChemiDoc MP imaging system (Biorad).

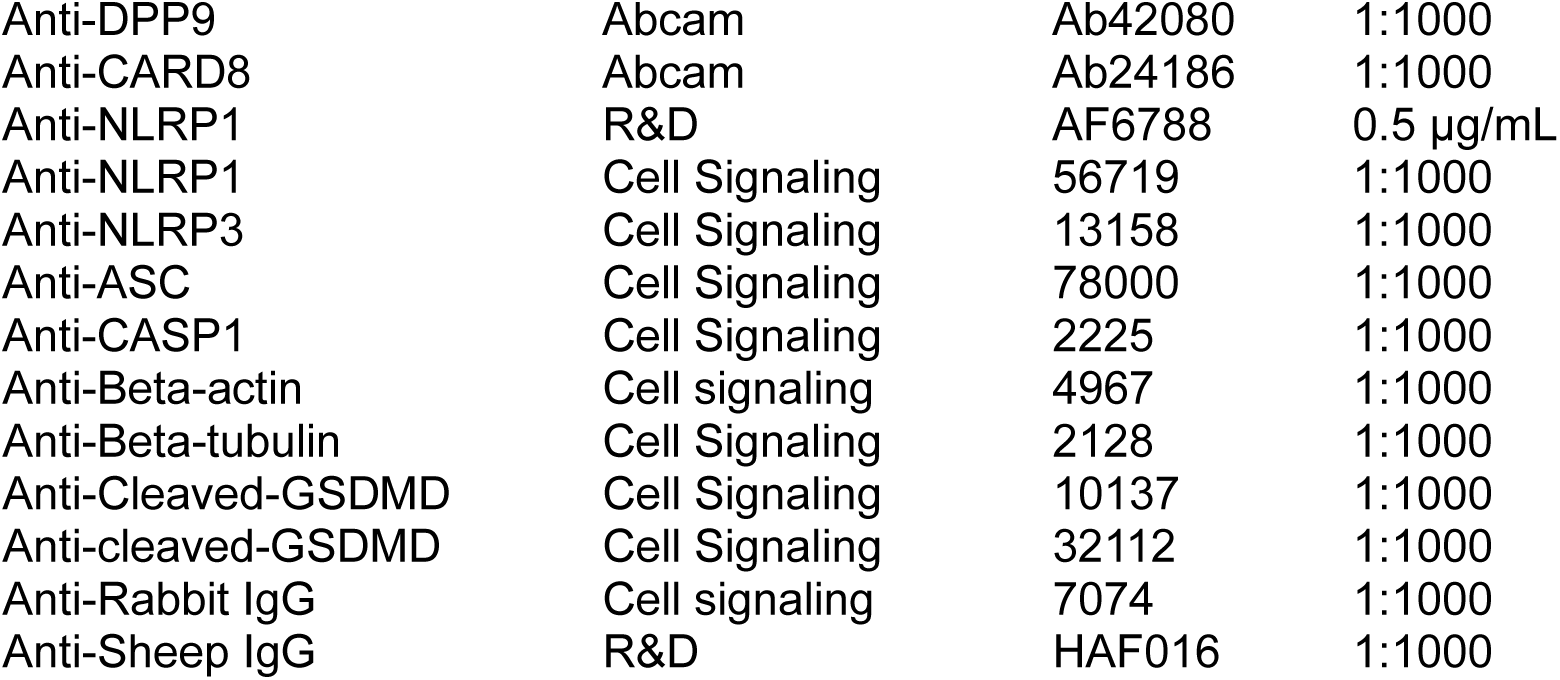

### Mice

*Dpp8* knock-out and *Dpp9* “floxed” mice were generated by the Flavell lab CRISPR Core as previously described^63^ using two sgRNAs that target exons containing the catalytic residues in C57BL6/N. Proper targeting was validated by sanger sequencing. *Dpp9* targeting vectors containing LoxP sequence were Ultramers ordered from IDT.

Genotyping primers and gRNA sequences:

mDpp8WTF: TCCCCCTCTTTGACAGGGTC

mDpp8WTR: CCATCAGGGAGAGGTAGCCA

139 bp WT band

mDpp8KOF: CCTCAGGGGTAATGTCCTCT

mDpp8KOR: GCTGACAGACTTGCTTCCAC

229 bp KO band

mDpp8 gRNA1: TGTTGCACTTAAGTGCTAAA

mDpp8 gRNA2: GCTCTGAGTGCTACTCTAGA

mDpp9F1: AGGGTGGTAGTGACTGAGAC

mDpp9R1: TGCTCTTAGACGCCAAGTAG

272 bp WT band

306 bp KI band

mDpp9 LoxP targeting vector 1: GGTGGTGATAATGGTGGTGGTGGTGATAGAGCTGATGGTGCCAATAACTTCGTATAATGTATGCTATACGAAGTTATGCCTGCACACATAAAGAACAGCTTTAGCAGGGCCCTCTGGGCA

mDPP9 gRNA1: TGTTCTTTATGTGTGCAGGC

mDpp9F2: GGACCCAGGCATCACTACAC

mDpp9R2: CAGGAAGGTGGGTACTCTCC

393 bp WT band

427 bp KI band

mDpp9 LoxP targeting vector 2: AACATTATGAAGCCACAAAAGCATCAAAGGCTATCACTTGCCTATAACTTCGTATAATGTATGCTATACGAAGTTATGAGCATCATGATTGTGCATAAGCATATAAAGTTCAAATAAAAT

mDPP9 gRNA2: TATGCACAATCATGATGCTC

Casp1 knock-out mice were generated by the Flavell lab and described previously^64^. Mutant mice for *Nlrp1*^44^ (JAX 021301)*, Il18*^65^ (JAX 004130)*, Il1r1*^66^ (JAX 003245) *and LysM-Cre*^46^ (JAX 004781) mice were purchased from the Jackson laboratories. *Nlrp1* mutant mice (JAX021301) were made by disrupting *Nlrp1b* on a 129 genetic background that lacks functional NLRP1a and NLRP1c and therefore lack all functional of all NLRP1 paralogues. Littermate control and experimental genotypes were weened together and cohoused while triple knock out mice were housed separately. For experiments using wild type controls, C57BL6/N mice bred in house were cohoused at least two weeks prior to experiment. Male and female mice 8-14 weeks of age were used unless otherwise noted. All animal experimentation was compliant with Yale Institutional Animal Care and Use Committee protocols. Statistical methods were not used to determine sample size, experiments were not randomized and investigators were not blinded unless otherwise stated in the methods. All mice generated for the purpose of this manuscript are publicly available upon request.

### Serum collection

Mice were euthanized with isoflurane and blood was collected from retroorbital bleeding, allowed to clot at room temperature and pelleted at 2,000G for 10 min. Serum was collected and stored at −80°C.

### LDH and Cytokine quantification

For ELISAs and LDH Release assays, 1-2 x 10^5^ THP1 or human/mouse monocytes were treated for 20-24 hours in 200 µL total volume. 2.5 x 10^4^ human macrophages or DCs were primed with 100 ng/mL LPS for 3 hours and stimulated for 20-24 in 200 µL total volume. All assays were conducted according to manufacturer’s guidelines. Mouse cytokine/chemokine multiplex ELISA (MD44) was conducted by Eve technologies. Graphs were generated and statistical analysis was conducted using GraphPad Prism 10.

LDH Release (Thermo EEA013)

Cytometric Bead Array for human IL-1β (BD 561509)

ELISA for human IL-18 (R&D DY318-05)

ELISA for mouse IL-1β (R&D DY401)

ELISA for mouse IL-18 (R&D DY7625, Invitrogen BMS618-3).

### Histology

Mice were euthanized with isoflurane and indicated organs were harvested, washed in PBS then fixed in 10% neutral buffered formalin overnight. Samples were then processed, paraffin embedded, sectioned and stained by the Yale Histology Core. Images were obtained on Keyence BZ-X microscope.

### Pathology Scoring

All histopathological assessment and scoring of H&E were performed on blinded samples by a board-certified pathologist as previously described^67,68^. Lung inflammation was scored semi quantitatively based on the abundance of intra-alveolar macrophages, infiltration of lung parenchyma by lymphocytes and macrophages, expansion of inter alveolar septa and development of granulomas.

### Single Cell RNA sequencing

In vivo labeling was used to discriminate between circulating and tissue resident immune cells. 2 μg of CD45-PERCP (Biolegend, 102129) was injected i.v. and after 10 minutes mice were euthanized using isoflurane. The lung was then inflated with 1 mL of digestion buffer A (1 mg/mL Elastase (Worthington LS002292), 500 μg/mL Dispase (Sigma D4693) and 10 μg/mL DNAse I (Roche 10104159001) in DMEM without serum), the trachea was tied shut and the lung was removed and incubated at 37°C for 45 minutes. The lung was then torn into small pieces and passed through a 100 μm strainer and the flow through-cell suspension was collected while the trachea was removed. This strainer was washed twice with 5 mL of serum free DMEM and once with 5 mL of DMEM supplemented with 10% fetal bovine serum and each was collected and pooled. The strainer containing undigested lung tissue was then placed into a 6 well dish containing digest buffer B (50 μg/mL liberase (Roche 05401127001) + 10 μg/mL DNAse I (Roche 10104159001) and digested for 37°C for 30 minutes. Cell suspension was collected and the cell strainer was washed twice with DMEM and pooled with previously collected cell suspensions. These were then pelleted and resuspended in 5 mL of ACK red blood cell lysis buffer, incubated at 5 minutes at room temperature, and washed in 5 mL of FACs buffer (PBS containing 1% fetal bovine serum and 1 mM EDTA). Cell suspension from 3 WT, 3 DKO and 1 DKO-severe mice were pooled by genotype, counted and stained with Anti-CD45.2-FITC (Biolegend, 109805) and 1 μM DAPI. 3 populations of cells were sorted for each sample namely: live immune cell (FITC-CD45.2^+^, Percp-i.v.CD45^-^, DAPI^-^), live non-immune cell (FITC-CD45.2^-^, Percp-i.v.CD45^-^, DAPI^-^) and auto-fluorescent^high^ alveolar macrophages. These 3 populations were mixed back for generating 10X library at 3:3:1 ratio to target a total 10,000 cells per sample in 0.04% BSA PBS. Library construction was completed by following the 10X Chromium Next GEM Single cell 3’ v3.1 protocol. With dimension reduction and clustering, we identified 9 major cell categories based on the expression of canonical markers, which includes epithelial cells (*Sftpc*, *S*ftpa1), fibroblasts (*Mgp*, *Mfap4*), endothelial cells (*Cldn5*, *Lyve1*), mesothelial cells (*Msln*, *Gpm6a*), T cells (*Trac*, *Cd3g*, *Cd3d*), B cells (*Igkc*, *Ighm)*, DCs (*Wdfy4*, *Irf8*), monocytes/macrophages (*Chil3*, *Lpl*) and neutrophils (*S100a8*, *S100a9*).

### scRNA-seq data analysis

The raw fastq output files were processed with CellRanger. The cellranger filtered matrix output directories were further read into R (4.2.0) using Read10X function from the Seurat package (5.1.0). Seurat Objects were created for each sample namely “WT”, “DKO” and “DKO-severe” and filtered for cells with 500 < nFeature_RNA < 5000, nCount_RNA > 1000 and mitochondria percentage < 10. 3 objects were merged into one for further processing including log normalization (scale.factor = 10,000), find variable features and scale data. Dimensionality reduction was performed by running “RunPCA” followed by “RunUMAP”. Cell clusters were identified unbiasedly using FindNeighbors and FindClusters. The markers of each cluster were identified by FindAllMarkers and cell identity was assigned to each cluster manually based on the existing knowledge. We also grouped similar Seurat clusters into cell categories and further subgroup them for zoom-in differential expression analysis with Seurat function FindMarkers and KEGG analysis. For the reproducibility of the analysis, the raw fastq files, cellranger filtered matrix outputs as well as the R Data for the Seurat objects generated by our study will be shared.

### Flow cytometry

Lungs were homogenized in gentleMACS Tubes with dissociator V1.02 (Miltenyi Biotec) in 5 mL of RPMI supplemented with 5% FBS, Penicillin, Streptomycin, Glutamine, 100 µg/mL Collagenase D (Sigma 11088882001) and 10 μg/mL DNAse I (Roche 10104159001). The tissue was then incubated at 37°C at 250 RPM for 30 min. Cell suspensions were pelleted, and resuspended in 5 mL of ACK lysis buffer for 5 minutes at room temperature, washed twice in FACS buffer and filtered through 100 micron cell strainers. Cells were then stained using antibody cocktail combined with anti-mouse CD16/32 (Bioxcell BE0307, clone 2.4G2) and fixable viability dye (Invitrogen, eFluor506) for 30 minutes at 4°C in the dark. Cells were then washed twice with FACS buffer, fixed in 4% paraformaldehyde for 10 minutes at room temperature.

For FLICA staining, cell suspensions were cultured with FLICA-660 (Antibodies inc, 9122) in RPMI supplemented with 10% fetal bovine serum, 100 units/mL penicillin, 100 µg/mL streptomycin and 2mM L-glutamine at 37°C for 1 hour prior to surface staining and fixation.

For cytokine staining cell suspensions were cultured in 100 µL of RPMI with 5% FBS, penicillin, streptomycin & glutamine supplemented with 25 ng/mL PMA (Sigma P1585) and 1µg/mL ionomycin (Sigma 10634) in a tissue culture incubator. After 1.5 hours 100 µL of stimulation media containing 2x GolgiPlug (BD 555029) was added to each sample and cultured an additional 1.5 hours. Cells were then pelleted and surface antigens were stained as described above. Cells were then fixed and permeabilized according to BD cytoplasmic kit (BD 555028) and cytokines were stained overnight at 4°C in the dark.

Stained and fixed cells were filtered, mixed with 10 µL of counting beads (Life technologies, AccuCheck) and analyzed on a BD LSRII. Flow cytometry data was analyzed with FlowJo_v10.8.1. Graphs were generated and statistical analysis was conducted using GraphPad Prism 7.

| Antigen | Fluorophore | Supplier | Catalogue | Clone |
| --- | --- | --- | --- | --- |
| CD45.2 | FITC | BD | 55372 | 104 |
| Siglec F | BV711 | BD | 740764 | E50-2440 |
| CD11c | PECy7 | Biolegend | 117318 | N418 |
| CD11b | AF700 | Biolegend | 101222 | M1/70 |
| Ly6G | APC | Biolegend | 127614 | IA8 |
| IAIE | APC-Cy7 | Biolegend | 107628 | M5/114.15.2 |
| XCR1 | BV421 | Biolegend | 148216 | ZET |
| SIRPα | PERCPCy5.5 | Biolegend | 144010 | P84 |
| Ly6C | BV605 | Biolegend | 128036 | HK1.4 |
| CD64 | PE | Biolegend | 139304 | X54-5/7.1 |
| NK1.1 | FITC | Biolegend | 108706 | PK136 |
| CD8a | BV711 | Biolegend | 100748 | 53-6.7 |
| TCRb | PECy7 | Biolegend | 109222 | H57-597 |
| CD4 | BV605 | Biolegend | 100548 | RM4-5 |
| CD44 | APC-Cy7 | Biolegend | 103028 | IM7 |
| CD62L | PERCPCy5.5 | Biolegend | 104432 | MEL-14 |
| CD45.2 | APC | Biolegend | 109814 | 104 |
| IFNγ | BV421 | Biolegend | 505830 | XMG1.2 |
| CD19 | PE | Biolegend | 115508 | 6D5 |
| CD45 | PERCPCy5.5 | Biolegend | 103132 | 30-F11 |
| Ly6G | FITC | Biolegend | 127606 | IA8 |

## Supporting information

Supplemental Tables (all)

## Code availability

All code will be made available upon request.

## Acknowledgements

The authors thank J. Alderman, C. Hughes, L. Evangelista, E. Hughes-Picard, B. Cadugan, C. Lieber, F. Zhang, P. Ranney, W. Philbrick and J. Horrocks for technical and administrative help. We would also like to thank L. Shan and E. Elliot for critical input on the manuscript. J.R.B. was supported by a Postdoctoral fellowship from the American Cancer Society. A.H. was supported by the China Scholarship Council. S.H. was supported by an Austrian Marshall Plan Foundation Master’s Fellowship. W.K.M. was supported by Cancer Research Institute/Irvington Postdoctoral Fellowship and a NIH T32-DK007356. A.H.N was funded by a Conquer Cancer – Pfizer Oncology Young Investigator Award. Any opinions, findings, and conclusions expressed in this material are those of the author(s) and do not necessarily reflect those of the American Society of Clinical Oncology® or Conquer Cancer®. This work was supported in part by NIH R21-AI178249 and HHMI (to R.A.F.).

## Author Contributions

J.R.B. conceptualized the project, designed and implemented experiments, analyzed data and constructed the manuscript. A.H. designed and implemented experiments, analyzed data including scRNA-seq analysis and contributed to conceptualization of the project and revision of the manuscript. A.H.N. conceptualized the human clinical research, curated and analyzed patient data and constructed the manuscript. E.B.F curated patient data. H.N.B., M.C., M.Z.M., C.Q. and N.G. facilitated collection of human primary immune cells. T.X. advised and conducted immune cell CRISPR/Cas9 experiments. H.M. scored murine lung and liver pathology. W.K.M. performed 10x scRNA-seq reactions. E.S. advised FLICA reporter experiments. S.H., M.C., T.I. and M.H.O. advised on immunophenotyping experiments. A.R. conducted ELISA experiments with mouse monocytes. E.A., M.M., T.K.C., M.R., M.T, M.J.S., M.R., and M.F. advised on clinical analysis of patient samples. F.F., J.A.S., D.C., D.G. and M.V.I. conducted and oversaw experiments on patient plasma conducted at UTSW. D.C. performed IL18 assay and associated analysis. M.V.I. collected the clinical data and oversees the IRB protocol. J.A.S. provided laboratory resources, intellectual input, and supervision. D.E.G. provided laboratory/biorepository resources, intellectual input, supervision. S.F., M.G. and K.S. conducted experiments on patient BAL. T.M. and C.J.Y. conducted eQTL analysis. A.G. conducted SNP imputation and provided patient data to A.H.N and E.B.F. R.A.F. provided resources, supervision and funding for this project, and contributed to conceptualization and tactics of the project and revision of the manuscript.

---

**Extended Data Figure 1.**
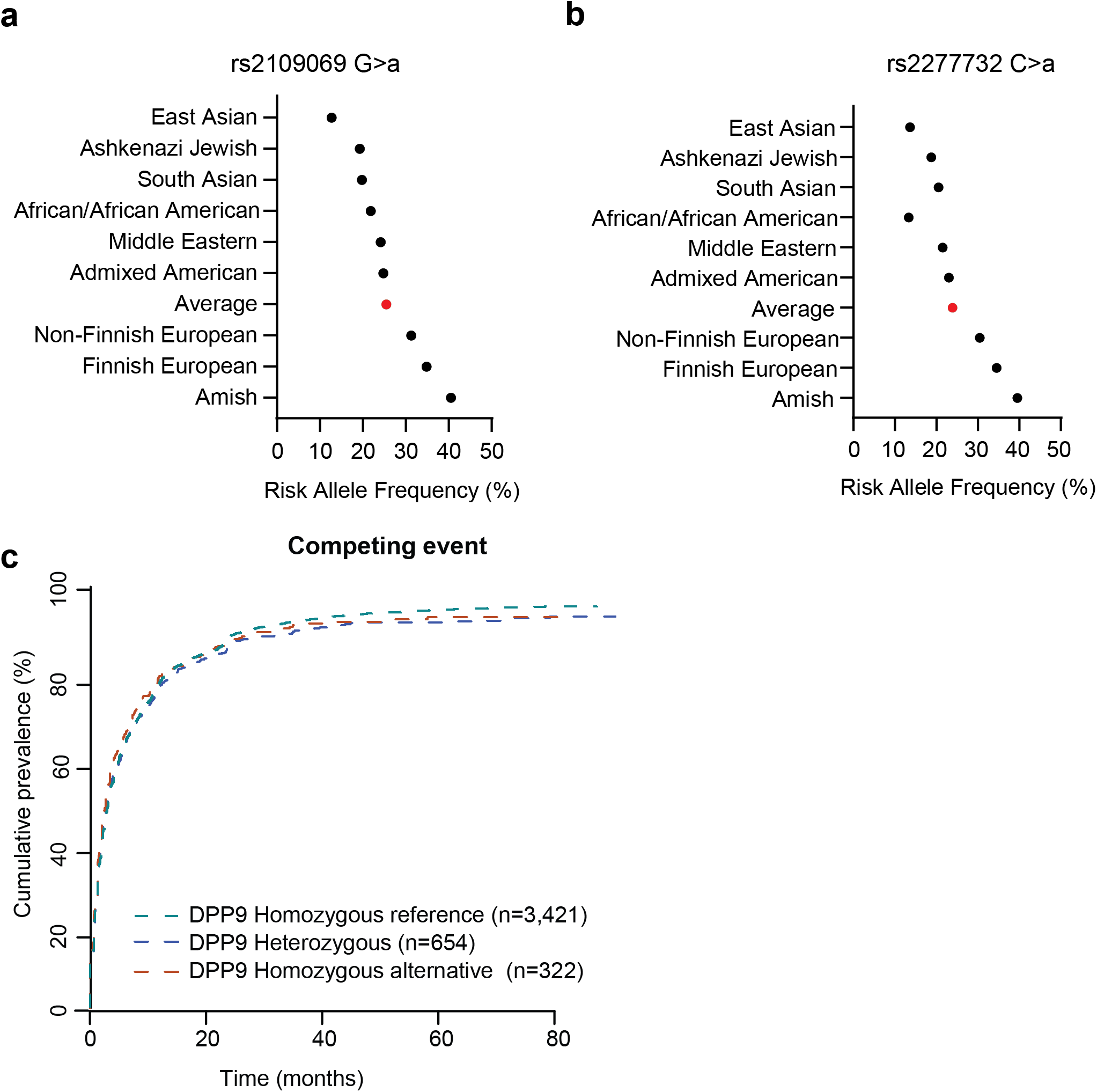
DPP9 risk allele frequencies and competing events. a-b) Risk allele frequency of *DPP9* disease associated SNPs rs2109069 and rs2277732 across nine ancestral populations, as reported in the Genome Aggregation Database version 4.1 (gnomAD v4.1). c) Cumulative incidence curves for non-CIP competing events resulting in ICI-discontinuation, estimated using the nonparametric Aalen–Johansen method. Patients in the DFCI cohort are stratified by SNP dosage. Red curves indicate patients homozygous for at least one of the three *DPP9* SNPs; blue curves represent heterozygous carriers; green curves depict patients lacking these risk alleles.

**Extended Data Figure 2.**
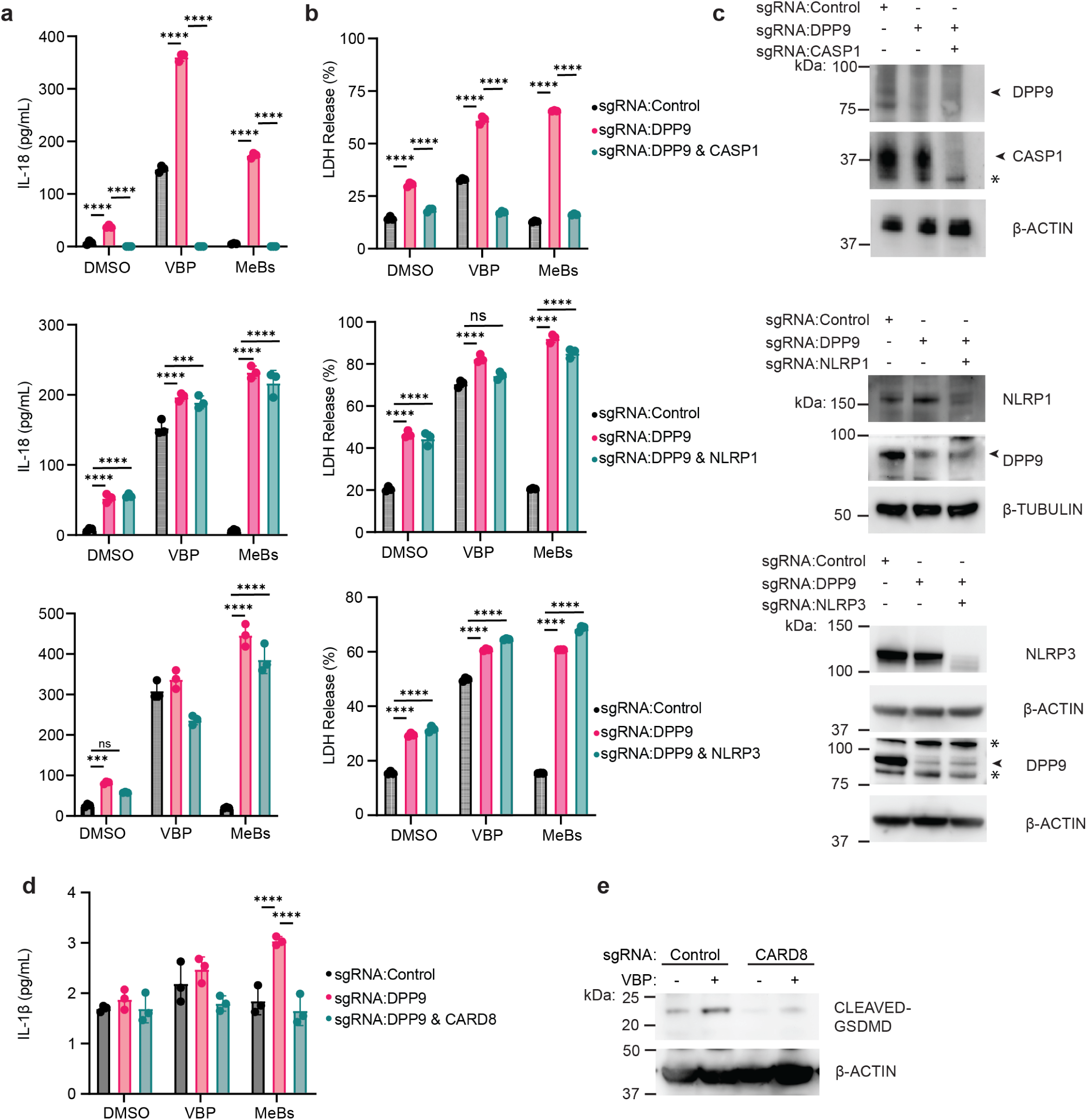
NLRP1 and NLRP3 are dispensable for VBP and MeBs responses in THP1 monocytes. THP1 monocytes expressing Cas9 were electroporated with indicated sgRNAs (*TRAC* was used as control). a) IL-18 ELISA, b) LDH release assay, c) immunoblot and d) IL-1β cytometric bead array. c) Immunoblot was used to confirm knock-out efficiency. Non-specific (*) and bands of interest (arrowhead) are indicated. e) Immunoblot of cleaved-GSDMD of indicated treatments. Experiments are representative of at least three independent experiments. Mean is shown with standard deviation of biological replicates corresponding to independently treated wells (n=3). Two-way ANOVA, Tukey’s multiple comparison test. \*\*\*\**p*<0.0001, \*\*\**p*<0.001. Valboro-pro (VBP) 25 µM, 10 µM methyl ester bestatin (MeBs) treated 24h.

**Extended Data Figure 3.**
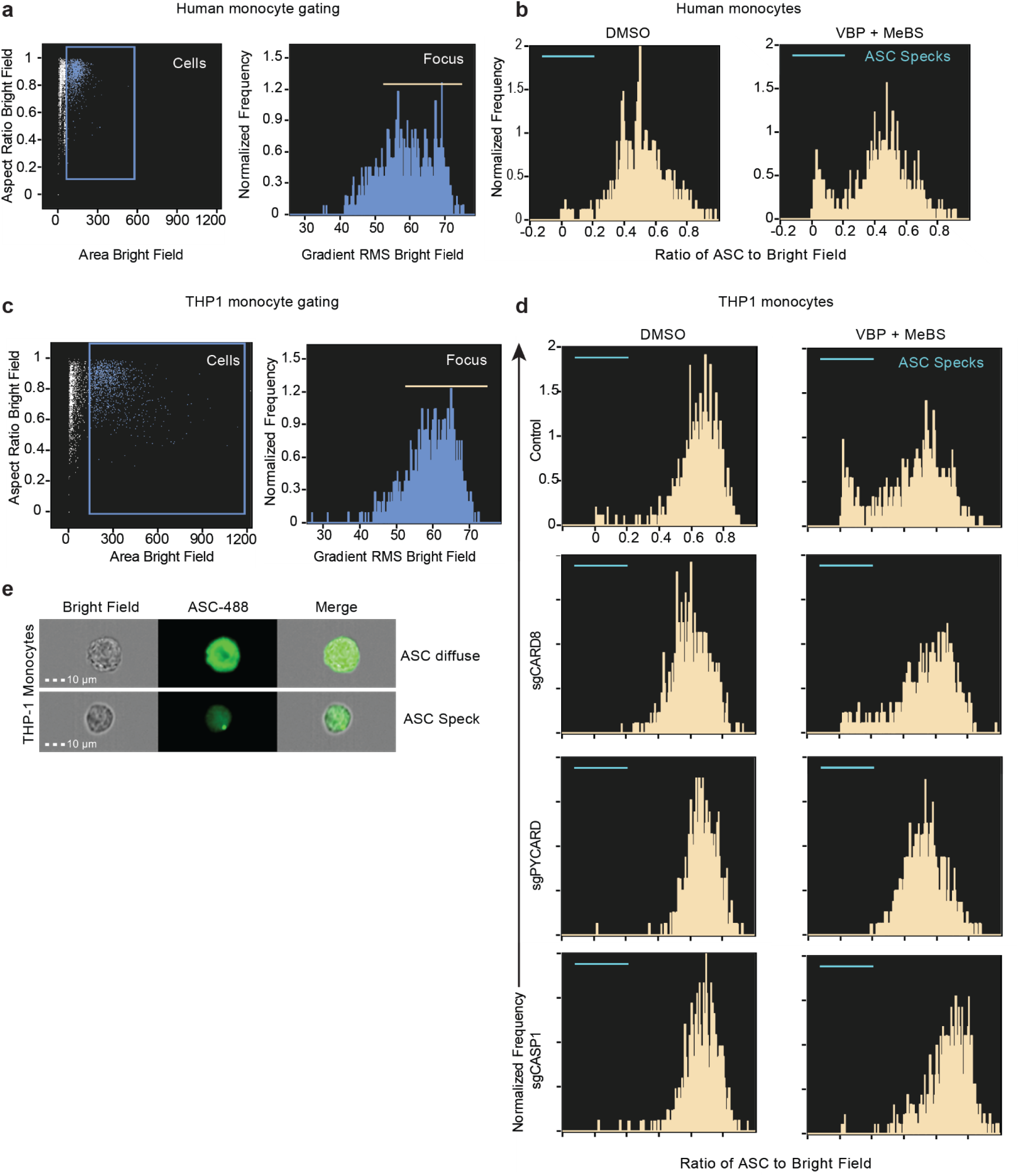
Imaging flow cytometry gating strategy. Gating strategy for primary human monocytes (a) and THP1 monocytes (c). Representative histograms of ASC “speck” quantification where cells with low ratio of ASC area relative to the size of the total cell (bright field) are calculated in primary human monocytes (b) and THP1 monocytes (d). THP1 cells expressing Cas9 were engineered by electroporating indicated sgRNAs (*TRAC* was used as control). treated 24h. f) Representative imaging cytometry data of ASC immunostaining (green) in THP1 monocytes. Diffuse and “speck” distribution represent inactive and activated/oligomerized state. Valbor-pro 25 µM (VBP), 10 µM methyl ester bestatin (MeBs).

**Extended Data Figure 4.**
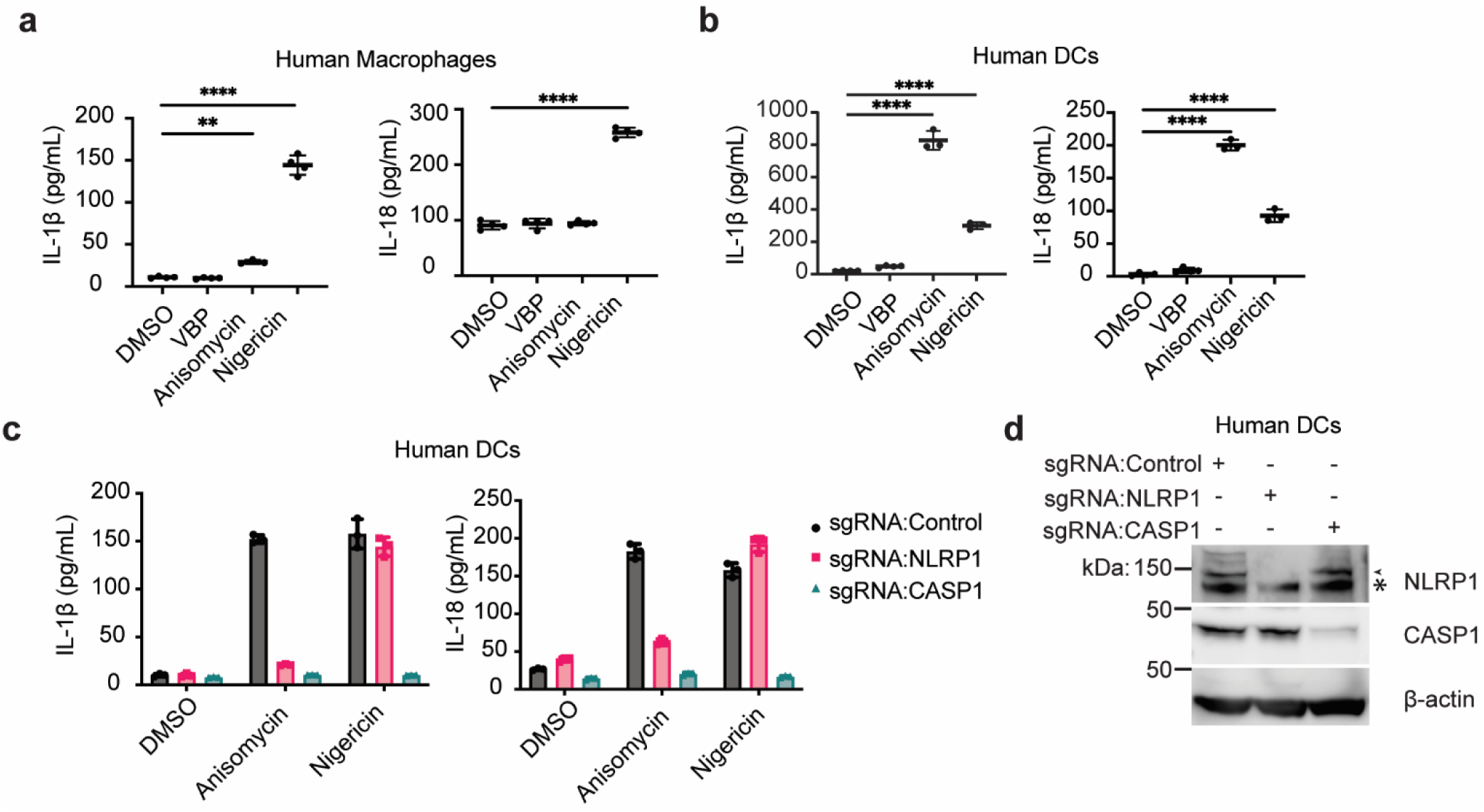
NLRP1 is activated in response to anisomycin in human dendritic cells (DCs). a-c) IL-1β and IL-18 ELISAs of human macrophage and DC supernatants. c) Primary DCs were engineered with Cas9 complexed with sgRNAs targeting Control (*TRAC*), *NLRP1* and *CASP1*. d) Immunoblot of DC protein lysate was used to confirm knock-out efficiency. Non-specific (*) and bands of interest (arrowhead) are indicated. Mean is shown with standard deviation of biological replicates corresponding to independently treated wells. Cells were primed with 100 ng/mL LPS 3 hours prior to addition of indicated treatments. Valboro-pro (VBP) 25 µM, 10 µM anisomycin, 10 µM Nigericin treated 24h. One-way ANOVA, Tukey’s multiple comparison test. \*\*\*\**p*<0.0001. \*\**p*<0.01. All results are representative of at least three independent experiments (n=3).

**Extended Data Figure 5.**
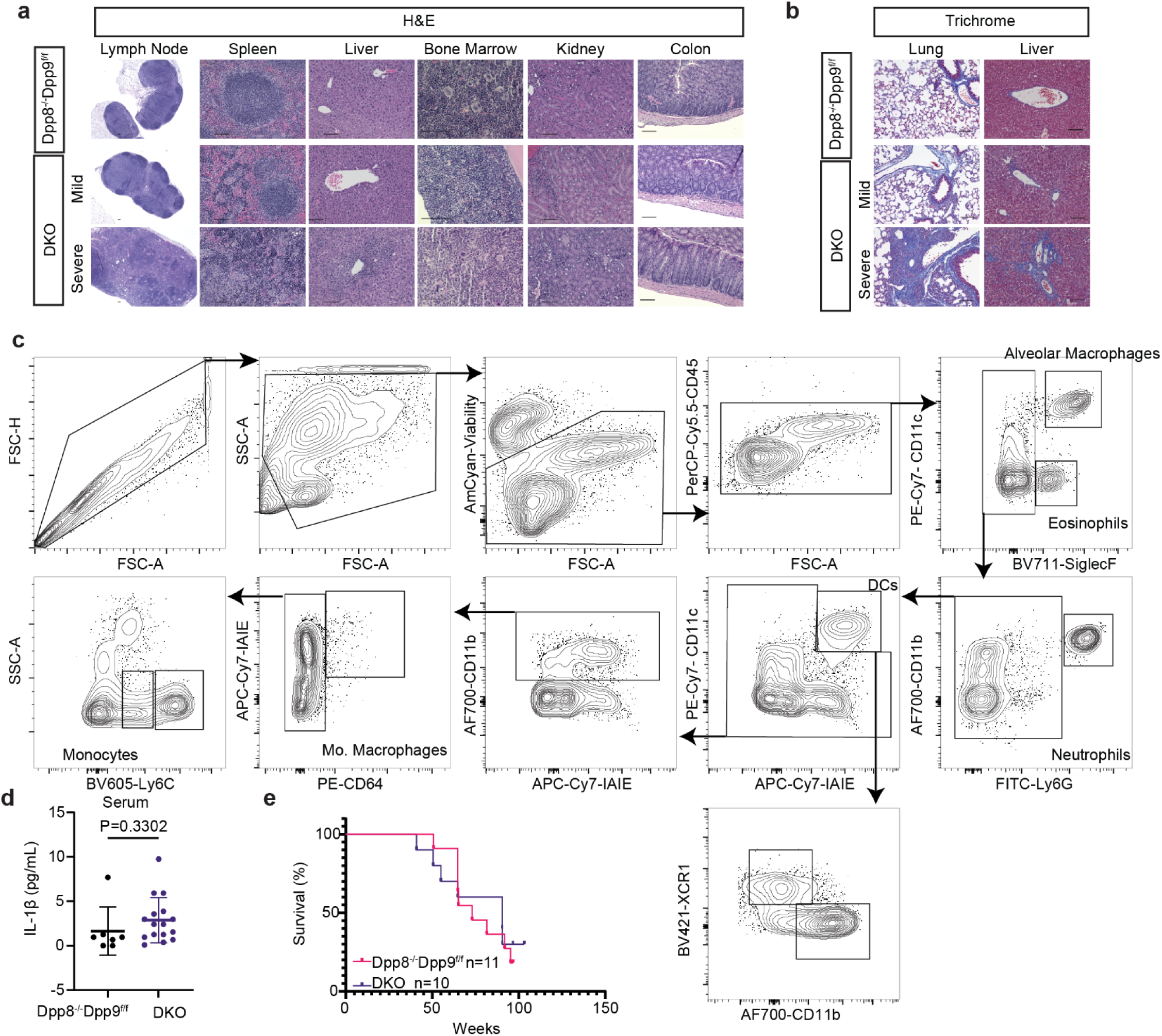
Histological analysis, FLICA gating strategy and survial. a-b) Hematoxylin and eosin (H&E, a) and trichrome (b) staining results of indicated organs. For unaffected organs, “severe” corresponds to the lung and liver severity of the mouse. At least 3 control, DKO-mild and DKO-severe mice were evaluated histologically for each organ. c) Gating strategy of lung myeloid cells from the FLICA CASP1 activity assay. Mo. (Monocyte). D) IL-1β ELISA from serum of mice with indicated genotypes (n=7-16). e) Kaplan-Meier survival curve of indicated genotypes. DKO, double knock-out (*Dpp8^-/-^Dpp9^f/f^LysM^Cre/+^*) TKO, triple knock-out.

**Extended Data Figure 6.**
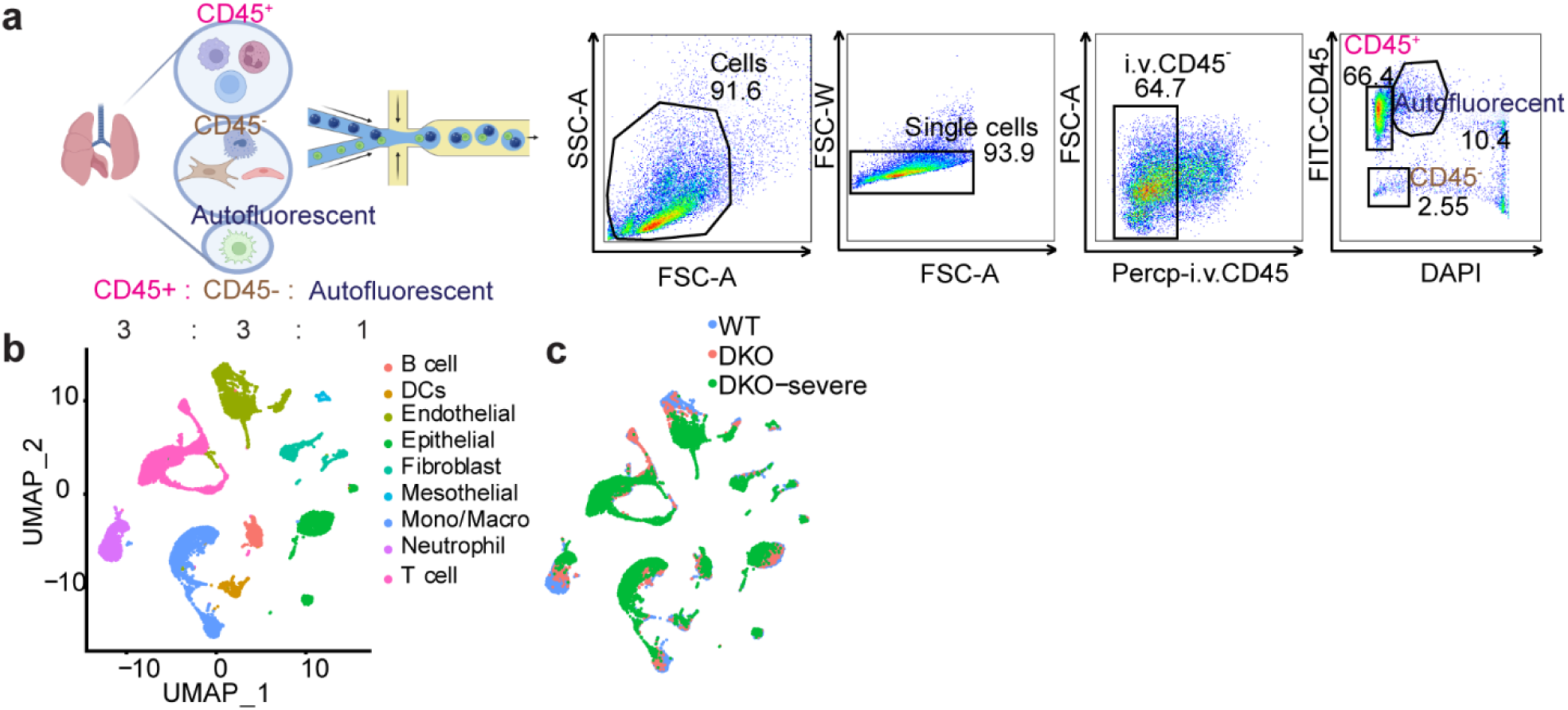
Single cell RNA sequencing strategy. a) Schematic illustration of scRNA-seq experimental design and gating strategy. CD45^+^, CD45^-^ and alveolar macrophages (auto fluorescent) were purified by cellular sorting and mixed together at a 3:3:1 ratio for sequencing using 10x platform (Biorender). scRNA-seq data from WT, DKO and DKO-severe shown as UMAP of b) cell clusters and c) indicated genotypes. DKO, double knock-out (*Dpp8^-/-^Dpp9^f/f^LysM^Cre/+^*).

**Extended Data Figure 7.**
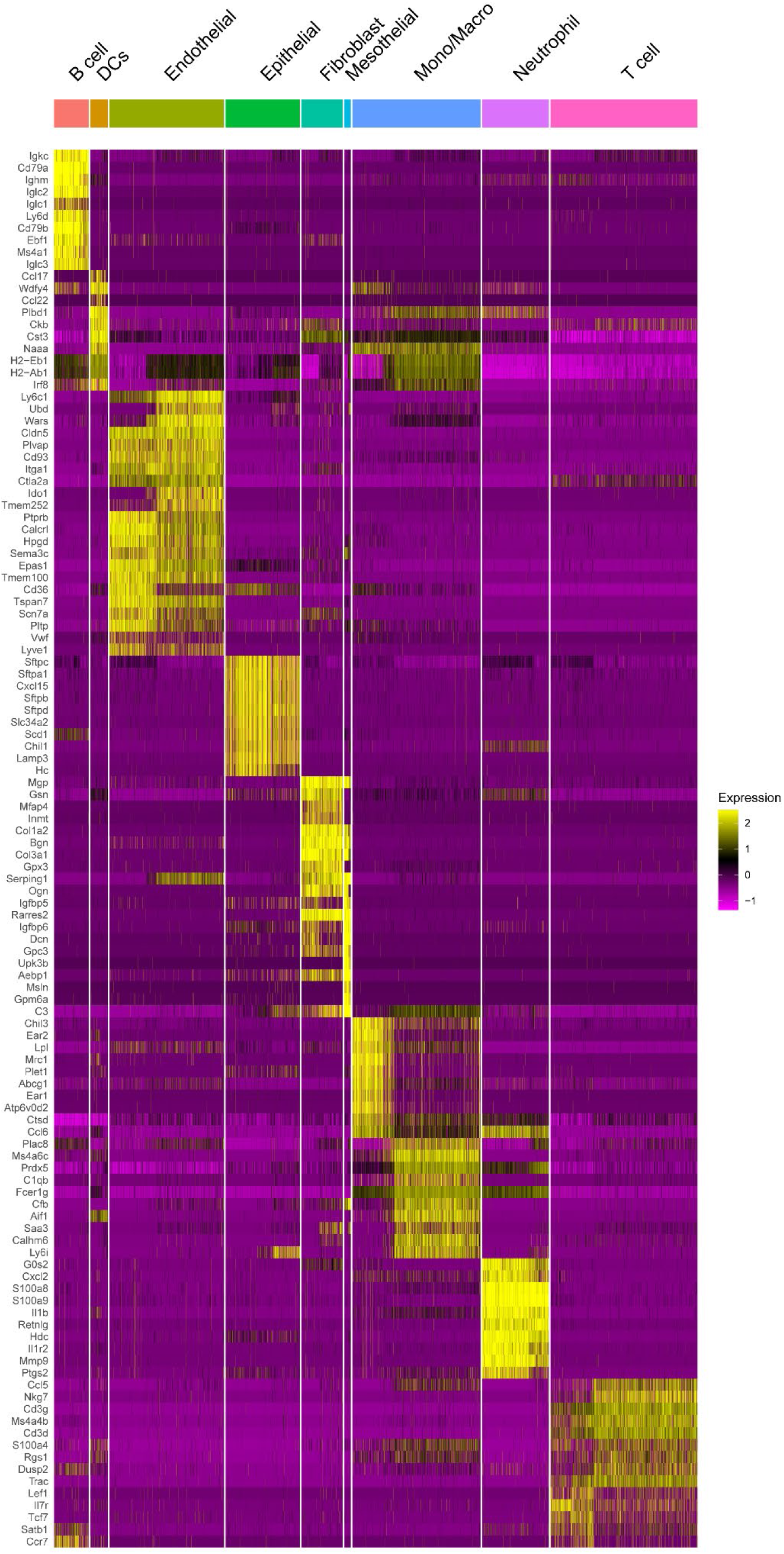
Heat map of lineage defining genes used to identify each cell type.

**Extended Data Figure 8.**
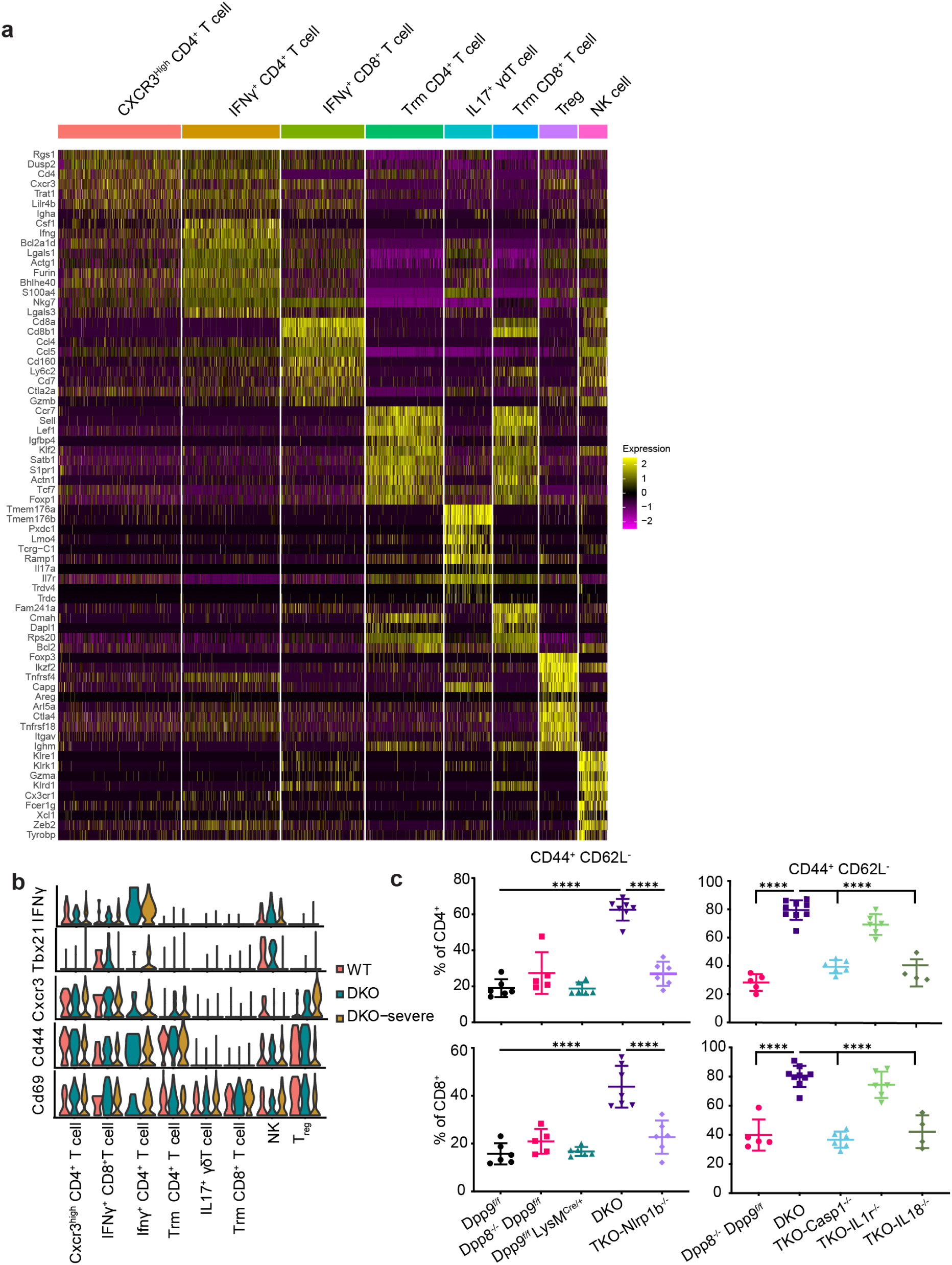
scRNA-seq and flow cytometry analysis of T cells. a) Heatmap highlighting cluster defining genes and cell type identities for indicated T cell subclusters. Tissue resident memory, (TRM). b) Violin plots of indicated T cell activation markers and genotypes. c) Flow cytometry quantification of CD44^+^ CD62L^-^ expression in CD4^+^ and CD8^+^ pulmonary T cells for indicated genotypes (n=4-9). One-way ANOVA, Tukey’s multiple comparison test. Mean is shown with standard deviation. Each dot represents values from a different mouse. \*\*\*\**p*<0.0001. Flow cytometry experiments are representative of at least two independent experiments. DKO, double knock-out (*Dpp8^-/-^Dpp9^f/f^LysM^Cre/+^*) TKO, triple knock-out.

**Extended Data Figure 9.**
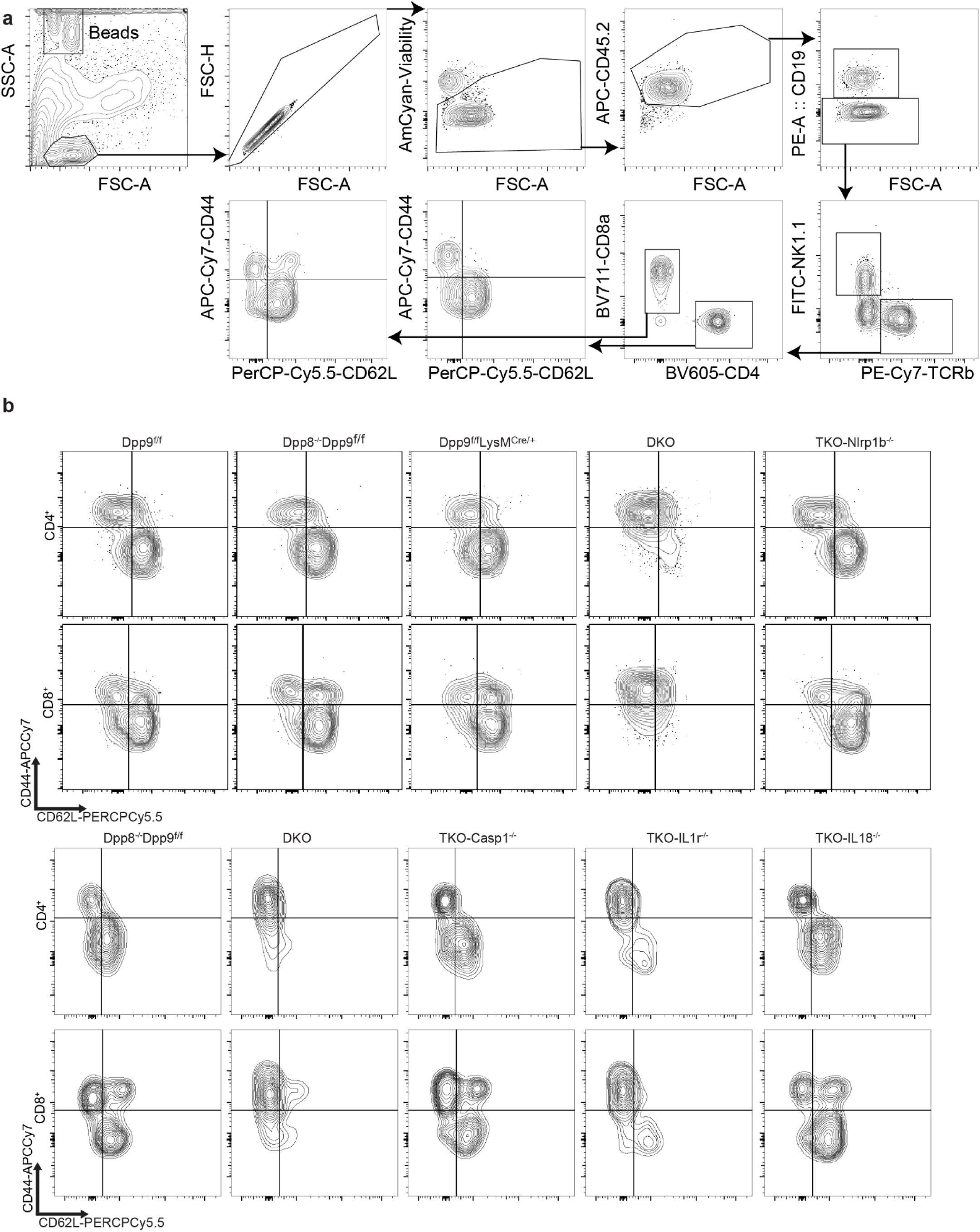
Gating strategy for T cell surface staining and representative flow plots. a) Gating strategy for surface staining of lung T cells. b) Representative flow panels showing CD44^+^ CD62L^-^staining in CD4^+^ and CD8^+^ pulmonary T cells of indicated genotypes. Quantification is shown in Extended Data Figure 6. DKO, double knock-out (*Dpp8^-/-^Dpp9^f/f^LysM^Cre/+^*) TKO, triple knock-out.

**Extended Data Figure 10.**
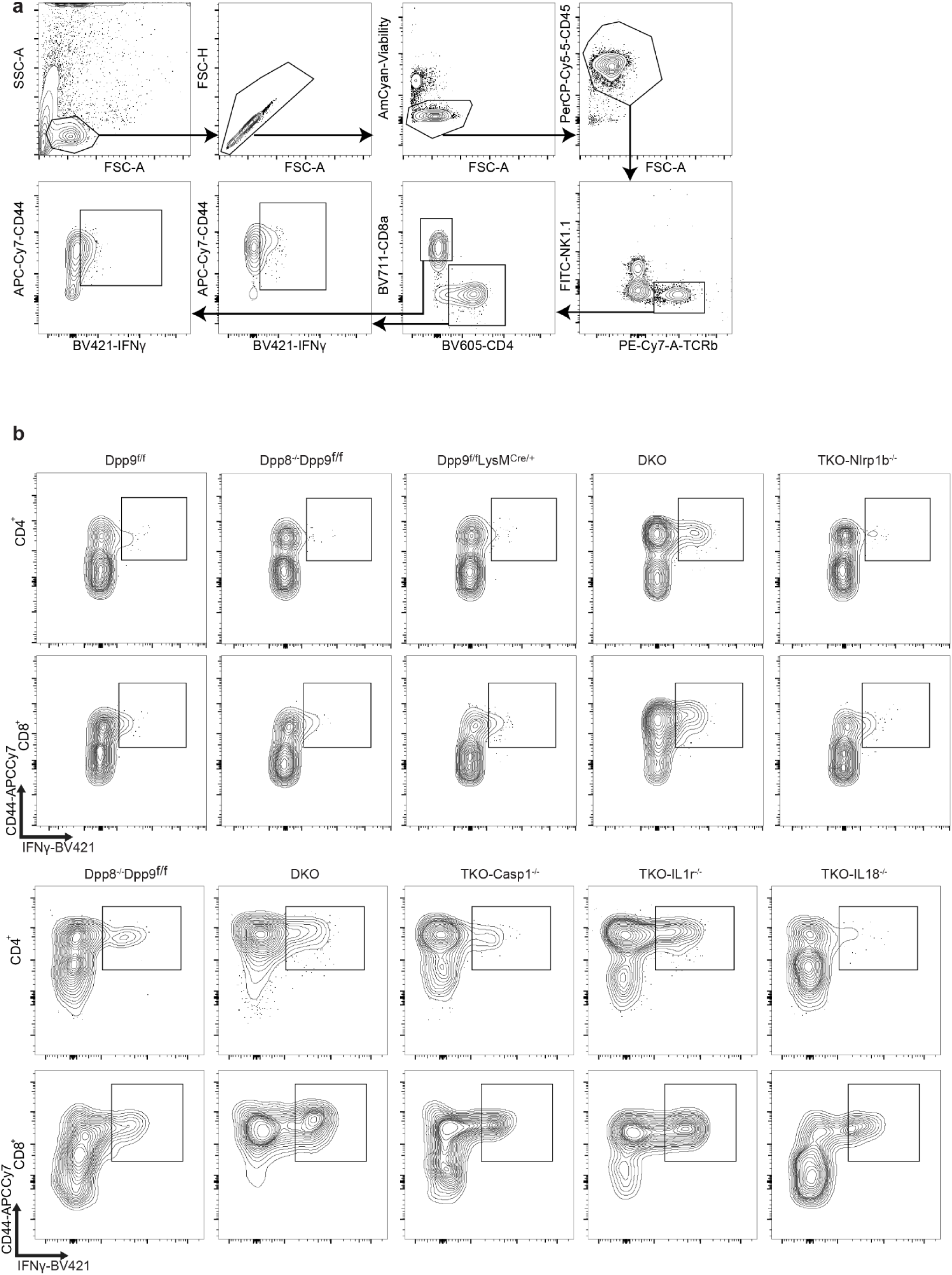
Gating strategy for T cell intracellular cytokine staining and representative flow plots. a) Gating strategy for intracellular cytokine staining of lung T cells. b) Representative flow panels showing CD44^+^ IFNγ^+^ CD4^+^ and CD8^+^ pulmonary T cells stimulated *ex vivo* with PMA and ionomycin of indicated genotypes. Quantification is shown in Figure 4. DKO, double knock-out (*Dpp8^-/-^Dpp9^f/f^LysM^Cre/+^*) TKO, triple knock-out.

**Extended Data Figure 11.**
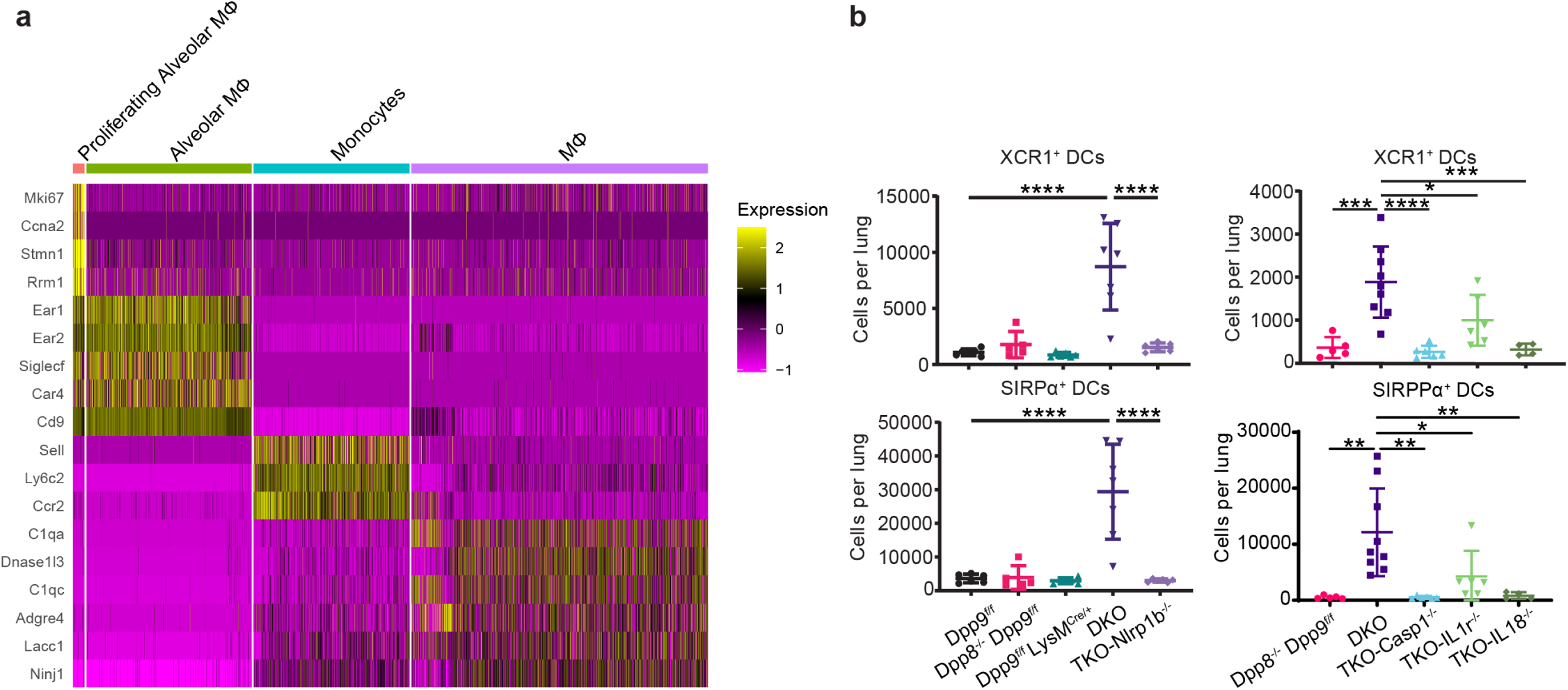
scRNA-seq and flow cytometry analysis of myeloid cells. a) Heatmap highlighting cluster defining genes and cell type identities for indicated myeloid subclusters. b) Flow cytometry quantification of dendritic cells (DCs) from indicated genotypes. For flow cytometry data, mean is shown with standard deviation. Each dot represents values from a different mouse (n=4-9). One-way ANOVA, Tukey’s multiple comparison test.\*\*\*\**p*<0.0001, \*\*\**p*<0.001, \*\**p*<0.01, \**p*<0.05. All flow cytometry results are representative of at least two independent experiments. DKO, double knock-out (*Dpp8^-/-^Dpp9^f/f^LysM^Cre/+^*) TKO, triple knock-out.

**Extended Data Figure 12.**
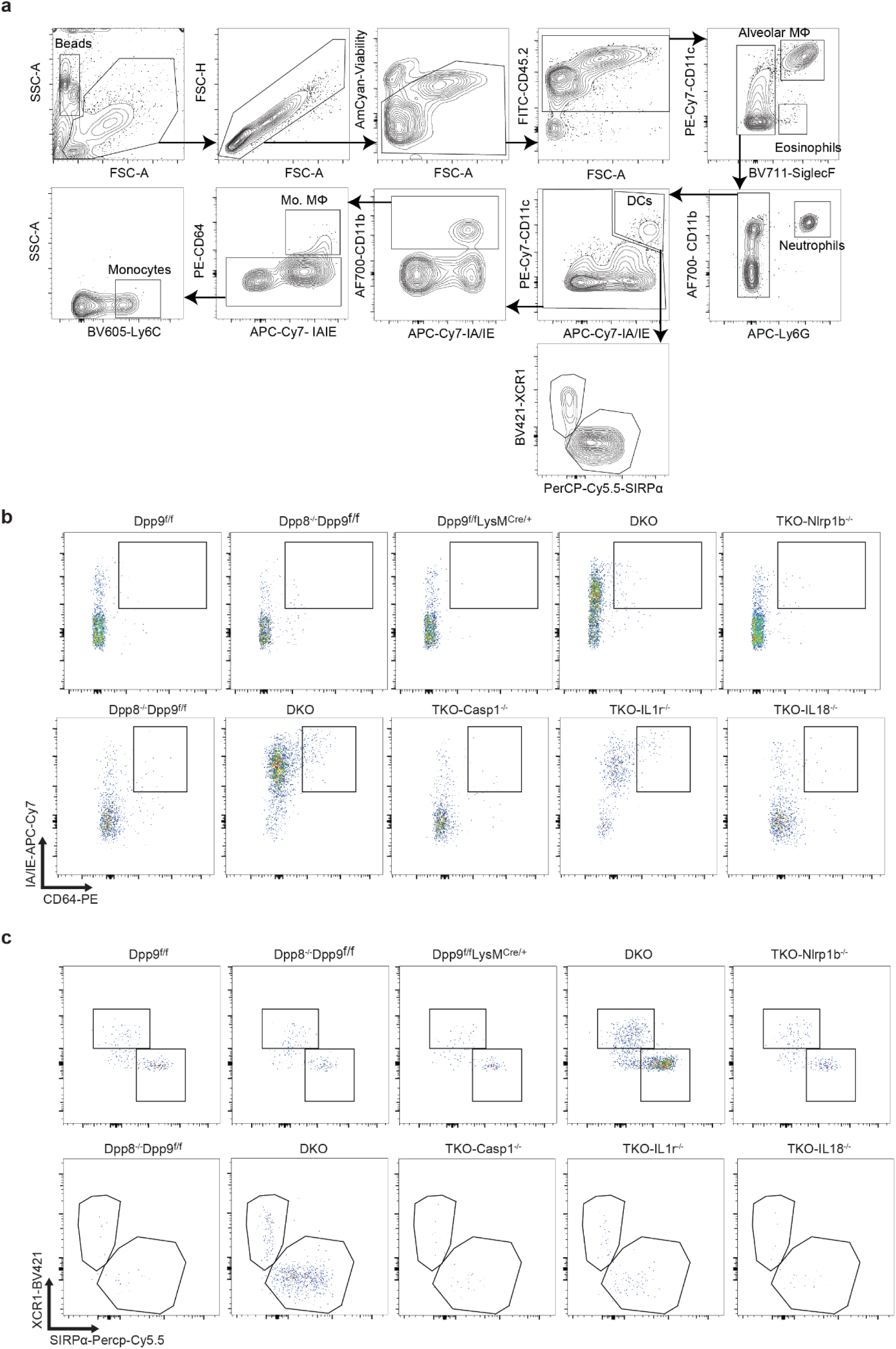
Gating strategy for myeloid cells and representative flow plots. a) Gating strategy for lung myeloid cells. b) representative flow panels showing b) monocyte derived macrophages indicating and c) dendritic cell (DC) cell populations. Quantification is shown in figure for and Extended Data Figure 8. DKO, double knock-out (*Dpp8^-/-^Dpp9^f/f^LysM^Cre/+^*) TKO, triple knock-out.

## Supplementary Tables

**Supplementary Table 1.** Clinical and demographic characteristics of the DFCI ICI-treated cohort. Summary of clinico-genomic variables for 4,397 cancer patients treated with immune checkpoint inhibitors at the Dana-Farber Cancer Institute, including age at ICI initiation, sex, race, genetic ancestry, cancer type, ICI regimen, and vital status at last follow-up. Values are reported as N (%) unless otherwise indicated.

**Supplementary Table 2.** Individual-level clinico-genomic data for the DFCI ICI-treated cohort.

**Supplementary Table 3.** Comparison of ICI-treated and non-ICI-treated patients in the DFCI cohort. Distribution of cancer type, vital status, sex, age at sequencing, and self-reported race among 4,397 ICI-treated patients and 15,473 non-ICI-treated patients who underwent tumor sequencing at DFCI.

**Supplementary Table 4.** Fine-Gray subdistribution hazard model for any-grade CIP in the full cohort. Multivariable competing-risks regression results for any-grade checkpoint inhibitor pneumonitis across the full DFCI cohort (n=4,397), modeling DPP9 genotype group (heterozygous and homozygous variant versus wild-type reference), age at ICI initiation, sex, and cancer category. Subdistribution hazard ratios (sHR), 95% confidence intervals, and p-values are reported.

**Supplementary Table 5.** Fine-Gray subdistribution hazard model for grade ≥3 CIP in the full cohort. Multivariable competing-risks regression results for grade ≥3 checkpoint inhibitor pneumonitis across the full DFCI cohort, modeling DPP9 genotype group (heterozygous and homozygous variant versus wild-type reference), age at ICI initiation, sex, and cancer category. Subdistribution hazard ratios (sHR), 95% confidence intervals, and p-values are reported.

**Supplementary Table 6.** Cause-specific Cox proportional hazards models for grade ≥3 CIP. Results of three separate cause-specific Cox models evaluating each DPP9 SNP (chr19:4723670_C / rs2277732, chr19:4717672_A / rs12610495, and chr19:4719443_G / rs2109069) as a continuous dosage score for grade ≥3 CIP risk, adjusted for age at ICI initiation, sex, cancer category, and top four genetic principal components (PC1–PC4). Hazard ratios (HR), 95% confidence intervals, and p-values are reported for each SNP model.

**Supplementary Table 7.** Cause-specific Cox proportional hazards models for grade ≥3 CIP in the European ancestry subset. Sensitivity analysis restricting to patients of inferred European genetic ancestry (n=4,168), evaluating each of the three DPP9 SNPs (rs12610495, rs2109069, rs2277732) as continuous dosage scores for grade ≥3 CIP risk, adjusted for age at ICI initiation, sex, and cancer category. Hazard ratios (HR), 95% confidence intervals, and p-values are reported, demonstrating concordance with full-cohort estimates.

**Supplementary Table 8.** Cause-specific Cox proportional hazards models for grade ≥3 immune-mediated hepatitis and colitis. Results of cause-specific Cox models evaluating each of the three DPP9 SNPs (rs12610495, rs2109069, rs2277732) for association with grade ≥3 ICI-related hepatitis (left) and grade ≥3 ICI-related colitis (right), adjusted for age at ICI initiation, sex, and cancer category. No significant associations were observed, supporting the organ specificity of the DPP9–CIP association. Hazard ratios (HR), 95% confidence intervals, and p-values are reported.

**Supplementary Table 9.** Serum IL-18 measurements in healthy controls and ICI-treated patients with and without checkpoint inhibitor pneumonitis — UT Southwestern cohort. Deidentified patient-level data for 76 ICI-treated individuals, including demographic information (age, sex, race, ethnicity), cancer type, ICI regimen, pneumonitis status and date, time from ICI initiation to pneumonitis, and serum IL-18 concentrations (pg/mL) and log₂-transformed values at baseline and after 6 weeks of ICI therapy.

**Supplementary Table 10.** Clinical characteristics of patients in the Johns Hopkins BAL IL-18 cohort. Deidentified patient-level data for 13 individuals, including study identifier, ICI treatment and CIP status (ICI+/CIP-, ICI+/CIP+) vital status, age, sex, and cancer type. This cohort was used to measure IL-18 protein levels in bronchoalveolar lavage fluid from ICI-treated patients with and without checkpoint inhibitor pneumonitis.

